# Phosphorylated Lamin A/C in the nuclear interior binds active enhancers associated with abnormal transcription in progeria

**DOI:** 10.1101/682260

**Authors:** Kohta Ikegami, Stefano Secchia, Omar Almakki, Jason D. Lieb, Ivan P. Moskowitz

## Abstract

*LMNA* encodes nuclear lamin A/C that tethers lamina-associated heterochromatin domains (LADs) to the nuclear periphery. Point mutations in *LMNA* cause degenerative disorders including the premature aging disorder Hutchinson-Gilford progeria, but the mechanisms are unknown. We report that Ser22-phosphorylated Lamin A/C (pS22-Lamin A/C) was localized to the interior of the nucleus in human fibroblasts throughout the cell cycle. pS22-Lamin A/C interacted with a specific subset of putative active enhancers, not LADs, primarily at locations co-bound by the transcriptional activator c-Jun. In progeria-patient fibroblasts, a subset of pS22-Lamin A/C-binding sites were lost whereas new pS22-Lamin A/C-binding sites emerged in normally quiescent loci. These new pS22-Lamin A/C-binding sites displayed increased histone acetylation and c-Jun binding, implying increased enhancer activity. The genes near these new binding sites, implicated in clinical components of progeria including carotid artery diseases, hypertension, and cardiomegaly, were upregulated in progeria. These results suggest that Lamin A/C regulates gene expression by direct enhancer binding in the nuclear interior. Disruption of the gene regulatory rather than LAD function of Lamin A/C presents a novel mechanism for disorders caused by *LMNA* mutations including progeria.

**HIGHLIGHTS:**

- pS22-Lamin A/C is present in the nuclear interior throughout interphase.
- pS22-Lamin A/C associates with active enhancers, not lamina-associated domains.
- pS22-Lamin A/C-genomic binding sites are co-bound by the transcriptional activator c-Jun.
- New pS22-Lamin A/C binding in progeria accompanies upregulation of disease-related genes.

## INTRODUCTION

Nuclear lamins polymerize to form the nuclear lamina, a protein meshwork that underlies the nuclear membrane and provides structural support to the nucleus (Gerace et al., 1978; Aebi et al., 1986; Goldman et al., 1986). There are two major nuclear lamin types, A-type and B-type (Dittmer and Misteli, 2011). A-type lamins (Lamin A and Lamin C) are specific to vertebrates, expressed in differentiated cells, and encoded by *LMNA* in humans (Dittmer and Misteli, 2011). Lamin A and Lamin C have different C-terminal tails (aa567-664) due to alternative splicing but are otherwise identical (Dechat et al., 2008). Point mutations in the *LMNA* gene cause a spectrum of human degenerative disorders including cardiomyopathies, muscular dystrophies, and the premature aging disorder Hutchinson-Gilford progeria (Worman et al., 2009). The molecular mechanisms underlying the disorders caused by *LMNA* mutations remain unclear.

Nuclear lamins including Lamin A/C interact with large heterochromatin domains called lamina-associated domains (LADs), which contain mostly transcriptionally inactive genes (Pickersgill et al., 2006; Guelen et al., 2008; Ikegami et al., 2010; Meuleman et al., 2012; Lund et al., 2014; van Steensel and Belmont, 2017). By interacting with LADs, nuclear lamins are implicated in the spatial organization of chromosomal regions at the nuclear envelope (van Steensel and Belmont, 2017). However, whether nuclear lamins play a direct role in transcriptional silencing of genes located at the nuclear periphery remains unclear. Artificially tethering genes to the nuclear periphery or inserting gene promoters into LADs does not always result in transcriptional repression (Finlan et al., 2008; Reddy et al., 2008; Leemans et al., 2019). In addition, many gene promoters in LADs remain inactive when their activities are examined outside of LADs (Leemans et al., 2019). While one study reported that tethering of Lamin A/C to gene promoters resulted in transcriptional downregulation (Lee et al., 2009), genetic depletion studies have shown that Lamin A/C and other lamins are not required for repression of genes within LADs (Kim et al., 2011; Amendola and van Steensel, 2015; Zheng et al., 2018). Thus, whether nuclear lamins including Lamin A/C have direct roles in transcriptional regulation has remained unclear.

Lamin A and Lamin C have been observed in the interior of the nucleus, in addition to its localization at the nuclear lamina (Dechat et al., 2010a). Initial descriptions engendered a model in which Lamin A/C in the nuclear interior represented a long-sought “nuclear scaffold” protein (Hozák et al., 1995; Barboro et al., 2002). However, subsequent studies demonstrated that Lamin A/C localized to the nuclear interior was soluble and highly mobile (Broers et al., 1999; Shimi et al., 2008), thus present as a non-polymerized form and not constituting a scaffold structure. The specific function of Lamin A/C in the nuclear interior has been difficult to ascertain, mainly due to a lack of understanding about how Lamin A/C is directed to the nuclear interior and technical challenges isolating nuclear-interior Lamin A/C.

Polymerization and depolymerization of nuclear lamins, required for nuclear envelope breakdown and the cell cycle, are regulated by phosphorylation of specific serine residues (Gerace and Blobel, 1980; Heald and McKeon, 1990; Peter et al., 1990; Ward and Kirschner, 1990). Ser22 (S22) and Ser392 (S392) of Lamin A/C are well characterized and are known as “mitotic sites” because they are phosphorylated during mitosis, leading to Lamin A/C depolymerization (Heald and McKeon, 1990; Peter et al., 1990; Ward and Kirschner, 1990). Early studies reported that phosphorylation of S22 and S392 begins at late G2 stage and is mediated by CDK1/Cyclin B, a kinase complex that promotes cell-cycle progression from G2 to mitosis (Heald and McKeon, 1990; Ward and Kirschner, 1990; Georgatos et al., 1997). More recently, S22 and S392 phosphorylation have been reported in the nuclear interior of interphase cells (Kochin et al., 2014), suggesting that S22/S392-phosphorylated, non-polymerized Lamin A/C may represent a nuclear-interior pool of Lamin A/C in interphase cells (Torvaldson et al., 2015). Separate studies proposed that S22 and S392 phosphorylation are increased upon changes in the mechanical environment of the cell and promote Lamin A/C disassembly and degradation (Swift et al., 2013; Buxboim et al., 2014). Therefore, Lamin A/C S22/S392 phosphorylation has been associated with mitotic nuclear lamina disassembly, but also with alternate cellular contexts in which its function remains unclear.

Hutchinson-Gilford progeria is a rare, fatal, childhood syndrome caused by heterozygous *LMNA* mutations and characterized by multiple organ system involvement, including the heart, vasculature, connective tissues, and bones (Gordon et al., 2003). Progeria patients invariably develop arteriosclerosis that ultimately causes death from myocardial infarction, heart failure, or stroke by the second decade of life (Merideth et al., 2008). The progeria mutations cause abnormal *LMNA* mRNA splicing, producing a mutant Lamin A protein called “progerin” that lacks a 50-aa sequence within the C-terminal tail domain (Eriksson et al., 2003). This 50-aa sequence includes a protease cleavage site used for removal of the farnesylated C-terminus during Lamin A maturation, and its deletion results in permanent farnesylation of progerin that promotes nuclear membrane association (Goldman et al., 2004). The C-terminal domain of Lamin A/C also includes binding sites for chromatin and DNA (Zastrow et al., 2004), and the C-terminal deletion in progerin results in reduced DNA and chromatin binding affinity *in vitro* (Bruston et al., 2010). Because Lamin A, Lamin C, and progerin interact with each other, progeria mutations have been postulated to function as dominant negative alleles, in which progerin may disrupt the normal functions of wild-type Lamin A/C (Gordon et al., 2014; Lee et al., 2016). The prevailing models for progerin’s pathological activity include altered function at LADs (Gordon et al., 2014). For example, progeria-patient cells exhibit disruption of LADs (McCord et al., 2012) and loss of nuclear-peripheral heterochromatin (Shumaker et al., 2006). The effect of progeria mutations on the function of nuclear-interior Lamin A/C has not been explored.

In this study, we report that S22-phosphorylated Lamin A/C (pS22-Lamin A/C) binds to genomic sites characteristic of active enhancers genome-wide in human fibroblasts. pS22-Lamin A/C-binding sites corresponded to a specific subset of accessible chromatin sites co-bound by the transcriptional activator c-Jun. In progeria-patient fibroblasts, a subset of pS22-Lamin A/C-binding sites were lost while new pS22-Lamin A/C-binding sites emerged at abnormal locations. Gains of pS22-Lamin A/C-binding sites in progeria were accompanied by increased c-Jun binding, increased H3K27 acetylation, and up-regulation of nearby genes relevant to progeria pathogenesis. In contrast, LAD alterations could not explain most gene expression alterations in progeria cell lines. We propose that Lamin A/C in the nuclear interior positively modulates enhancer activity, separate from its role at LADs, and that Lamin A/C’s role at enhancers contributes to progeria pathogenesis.

## RESULTS

### Phospho-S22-Lamin A/C is localized to the interior of the nucleus throughout the cell cycle

We investigated pS22-Lamin A/C as a candidate functional non-polymerized Lamin A/C in the nuclear interior following reports that pS22-Lamin A/C represents a depolymerized form of Lamin A/C during mitosis (Gerace and Blobel, 1980; Heald and McKeon, 1990; Peter et al., 1990; Ward and Kirschner, 1990) and that pS22-LMNA is detectable in interphase cells (Kochin et al., 2014). We confirmed the specificity of an anti-pS22-Lamin A/C monoclonal antibody (antigen: synthetic phosphopeptide surrounding S22; **Fig. S1A**) for the S22-phosphorylated state using ELISA on synthetic Lamin A/C peptides (aa 2-30) with either phospho-S22 or non-phospho-S22 (**Fig. S1B**). We also identified a reference anti-pan-N-terminal-Lamin A/C monoclonal antibody (antigen: synthetic non-phosphopeptide aa2-29; **Fig. S1A**) that recognized both the phospho-S22 and non-phospho-S22 peptides, with approximately 4-fold higher reactivity toward the non-pS22 peptide (**Fig. S1B**). Immunofluorescence microscopy of TERT-immortalized human BJ-5ta fibroblasts using these antibodies revealed pS22-Lamin A/C signals localized to the nuclear interior but not at the nuclear periphery, whereas pan-N-terminal-Lamin A/C signals localized predominantly to the nuclear periphery with weak signals in the nuclear interior (**Fig. 1A**). The pS22-Lamin A/C and pan-N-terminal-Lamin A/C immunofluorescence signals were absent in BJ-5ta-derived *LMNA^−/−^* cells, confirming the specificity of the Lamin A/C signals (**Fig. 1A**). During interphase, pS22-Lamin A/C was detectable in the nuclear interior but not at the nuclear periphery while pan-N-terminal-Lamin A/C signals were observed predominantly at the nuclear periphery (*Early G1* and *Interphase* in **Fig. 1B**). At late G2, pS22-Lamin A/C signals increased dramatically, consistent with the model that S22 is highly phosphorylated during the G2-to-M transition for Lamin A/C depolymerization (Peter et al., 1990) (*Late G2* in **Fig. 1B**). During mitosis, when the nuclear envelope is absent, strong pS22-Lamin A/C signal was observed throughout the cytoplasm (*Prophase* and *Metaphase* in **Fig. 1B**). Interestingly, the anti-pan-N-terminal-Lamin A/C antibody did not produce signals during mitosis (**Fig. 1B**), suggesting that other mitotic modifications, such as phosphorylation at Ser12, Ser18, or Thr19 (Dephoure et al., 2008), may have affected the reactivity of this antibody during mitosis. Flow cytometry of BJ-5ta cells using the anti-pS22-Lamin A/C antibody confirmed that pS22-Lamin A/C was present in G0/G1, S, and G2/M phases (**Fig. 1C; Fig. S1C)**. Western blotting of cell-cycle synchronized BJ-5ta cells further confirmed persistent S22-phosphorylated Lamin A/C throughout the cell cycle, with apparently stronger pS22-Lamin C signals than pS22-Lamin A signals (**Fig. 1D, E; Fig. S1D, E**). Thus, in human fibroblasts, pS22-Lamin A/C exists in the nuclear interior throughout the cell cycle.

**Figure 1.**
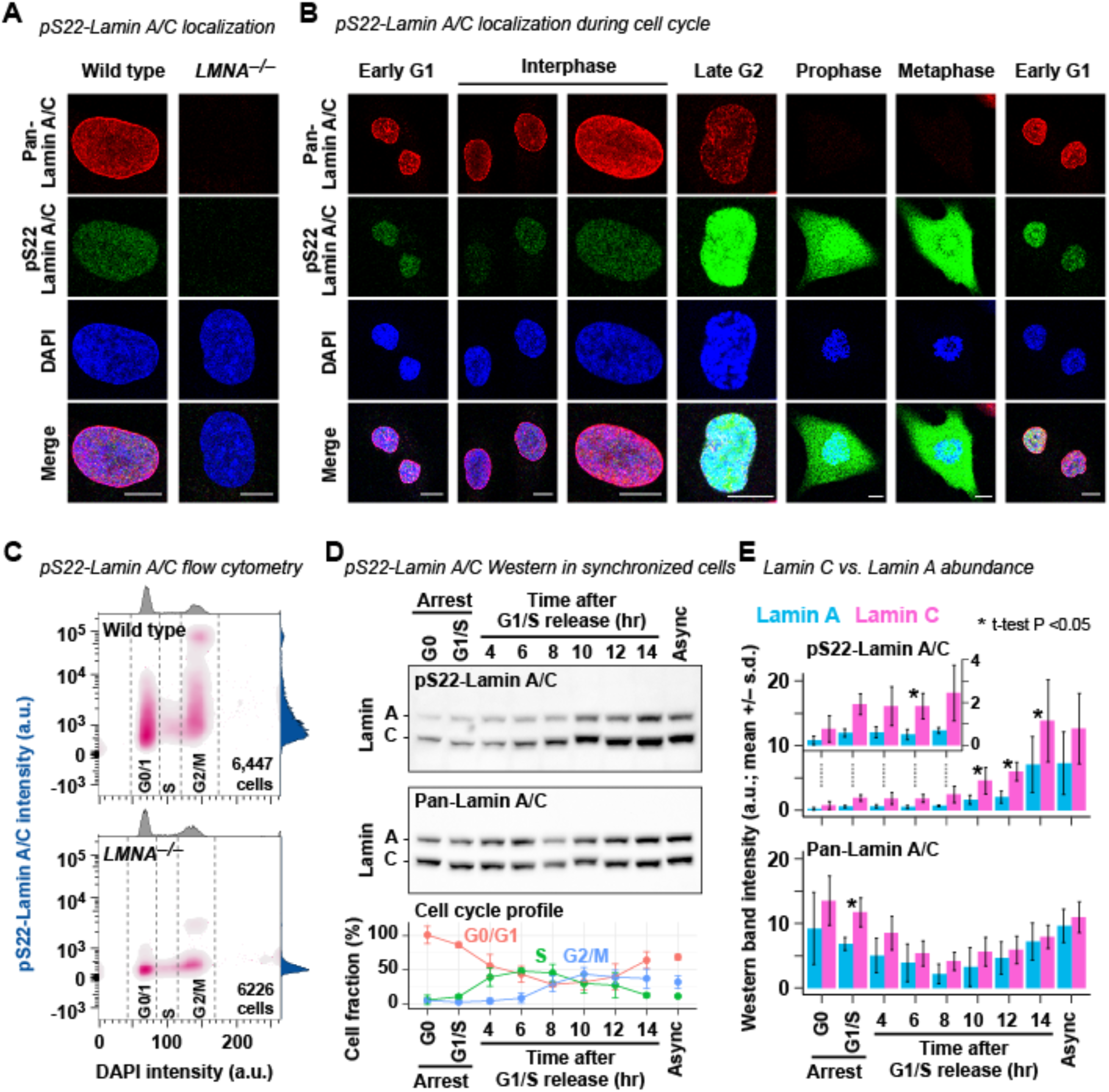
pS22-Lamin A/C is localized in the interior of the nucleus throughout the cell cycle. **(A)** Immunofluorescence using anti-pan-N-terminal-Lamin A/C (pan-Lamin A/C) and anti-phospho-S22-Lamin A/C (pS22-Lamin A/C) antibodies. Wild-type, TERT-immortalized human fibroblast BJ-5ta. *LMNA^−/−^*, BJ-5ta-derived *LMNA^−/−^* fibroblasts. Bar, 10 µm. See **Fig. S1A, B** for antibody validation. Representative images from 3 biological replicates. **(B)** Same as **(A)**, but BJ-5ta cells at specific cell-cycle stages are shown. **(C)** Flow cytometry of cells stained for pS22-Lamin A/C and DNA (DAPI), shown as two-dimensional density of cells. One-dimensional density plots are shown along axes. a.u., arbitrary unit. Representative analysis of 3 biological replicates. See **Fig. S1C** for no antibody control. **(D)** (Top) Western blot of cell-cycle synchronized BJ-5ta using anti-pS22-Lamin A/C and anti-pan-N-terminal-Lamin A/C antibodies. (Bottom) Cell-cycle stage fraction at each time point (mean +/– standard deviation) determined by flow cytometry. See **Fig. S1D, E** for uncropped images and biological replicates. **(E)** Quantification of Lamin A and Lamin C Western blot band intensities (mean of 3 biological replicates +/– standard deviation). Inset, narrower y-axis range for the first 5 time points. P, one sample *t*-test assessing distribution of log_2_[Lamin C]/[Lamin A] signal ratios for each time point.

### Phospho-S22-Lamin A/C interacts with genomic sites outside of lamina-associated domains

We hypothesized that pS22-Lamin A/C interacts with the genome because Lamin A/C has been shown to bind DNA and chromatin *in vivo* and *in vitro* (Taniura et al., 1995; Stierlé et al., 2003; Meuleman et al., 2012; Lund et al., 2014). To test this hypothesis, we performed ChIP-seq in asynchronous BJ-5ta cells using the anti-pS22-Lamin A/C antibody and using the anti-pan-N-terminal-Lamin A/C antibody as a comparison. We observed that pS22-Lamin A/C exhibited point-source enrichment at discrete sites located outside of lamina-associated domains (LADs) and not with LADs themselves, in sharp contrast to pan-N-terminal-Lamin A/C, which were strongly enriched at LADs (Guelen et al., 2008) (**Fig. 2A–C; Fig. S2A–D; Table S5,6**). We identified 22,966 genomic sites bound by pS22-Lamin A/C genome-wide. The pS22-Lamin A/C ChIP-seq signals were abolished in the BJ-5ta-derived *LMNA^−/−^* cell line, confirming the specificity of the pS22-Lamin A/C ChIP-seq signals in wild-type BJ-5ta (**Fig. 2B, D**). Pan-N-terminal-Lamin A/C ChIP-seq also detected weak signals at pS22-Lamin A/C-binding sites (**Fig. 2B, D**). To examine the chromatin localization of pS22-Lamin A/C with a different approach, we expressed Lamin A or Lamin C either with phospho-mimetic mutations (S22D/S392D), phospho-deficient mutations (S22A/S392A), or without mutations in *LMNA^−/−^* cells. We then performed immunofluorescence and ChIP-seq in these cells using the anti-full-length-Lamin A/C antibody. These studies revealed that phospho-mimetic Lamin C (S22D/S392D) was highly abundant in the nuclear interior (**Fig. S2E, F**) and strongly bound to pS22-Lamin A/C-binding sites (**Fig. 2E, F; Fig. S2H**). Phospho-mimetic Lamin C was still weakly localized at the nuclear periphery (**Fig. S2E**) unlike endogenous pS22-Lamin A/C (**Fig. 1A**), and correspondingly, was also enriched at LADs (**Fig. S2G, I**). In contrast, wild-type Lamin A and phospho-deficient Lamin A were strongly localized at the nuclear periphery (**Fig. S2E, F**) and strongly enriched at LADs (**Fig. S2G, I**), but showed no enrichment at pS22-Lamin A/C-binding sites (**Fig. 2E, F; Fig. S2H**).

**Figure 2.**
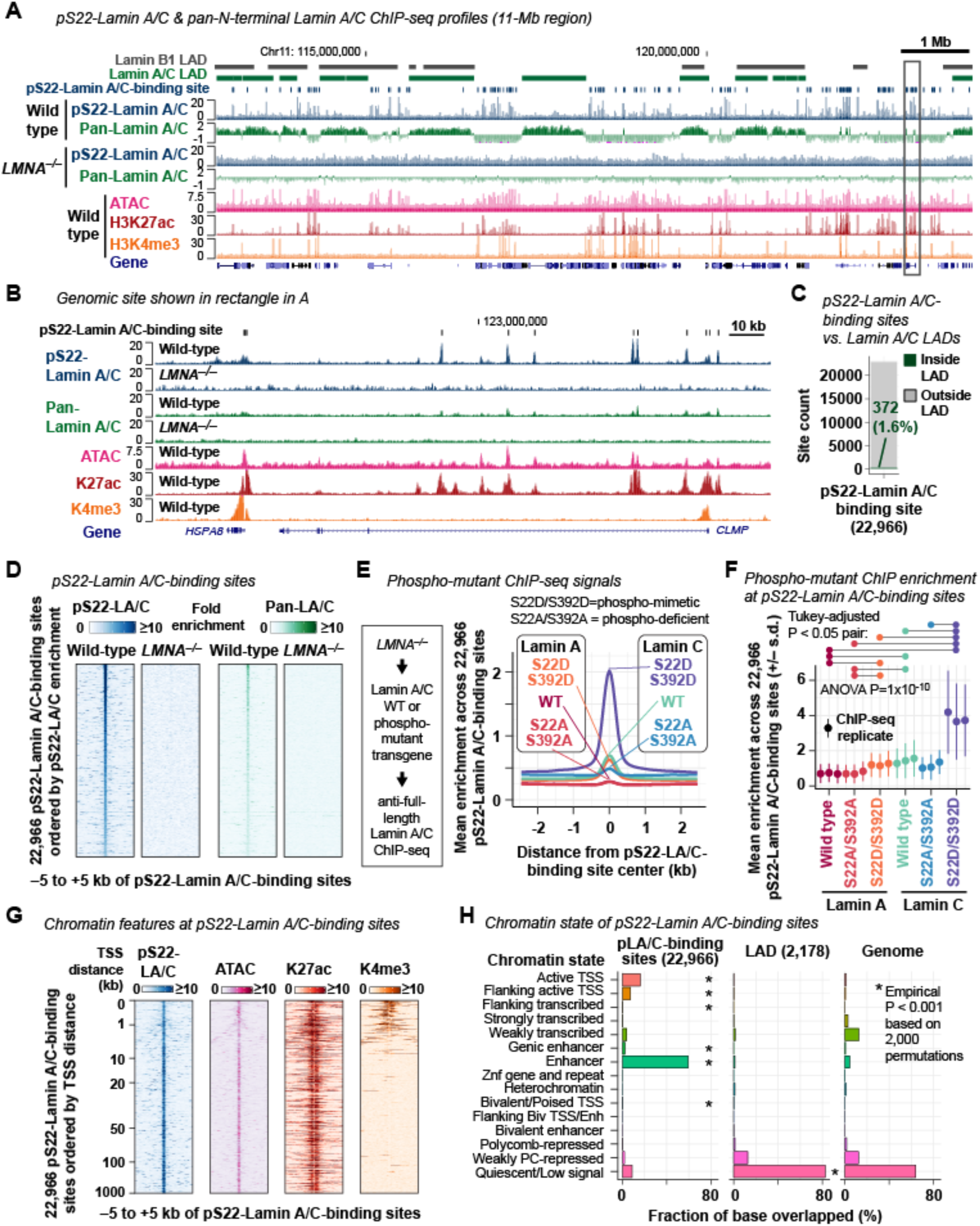
pS22-Lamin A/C associates with putative enhancers genome-wide. **(A)** Representative pS22-Lamin A/C ChIP-seq and pan-N-terminal-Lamin A/C ChIP-seq profiles. Histone ChIP-seq and ATAC-seq are shown for comparison. Signals are fold-enrichment (FE) scores, which are replicate-combined, depth- and input-normalized read coverage. Pan-N-terminal-Lamin A/C profiles are in the log_2_ scale to visualize LAD pattern. Lamin B1 LADs are defined by DamID in lung fibroblast (Guelen et al., 2008). Lamin A/C LADs are defined by pan-N-terminal-Lamin A/C ChIP-seq in BJ-5ta (this study). Wild type, BJ-5ta fibroblasts. *LMNA^−/−^*, BJ-5ta-derived *LMNA–/–* cells. All performed in ≥2 biological replicates (**Table S3**). See **Fig. S2A–D** for additional analyses on Lamin A/C ChIP-seq. **(B)** Region shown in rectangle in **(A)**. Pan-N-terminal-Lamin A/C ChIP-seq profiles are in the linear scale for a direct comparison with pS22-Lamin A/C profiles. **(C)** Positions of pS22-Lamin A/C-binding sites with respect to 2,178 Lamin A/C LADs. **(D)** pS22-Lamin A/C and pan-N-terminal-Lamin A/C ChIP-seq FE scores at pS22-Lamin A/C-binding sites. **(E)** ChIP-seq FE scores for transgene-driven Lamin A or Lamin C with phospho-deficient S22A/S392 mutations, or phospho-mimetic S22D/S392D mutations, or without mutation, expressed in *LMNA^−/−^*cells. ChIP was performed with anti-full-length-Lamin A/C antibody in 3 biological replicates. See **Fig. S2E-I** for protein localization and additional ChIP analyses. **(F)** ChIP-seq FE scores at pS22-Lamin A/C-binding sites for each biological replicate of **(E)**. Sum of FE scores within +/–250 bp of site center is averaged. One-way ANOVA compares means of all isoforms, with post-hoc Tukey analysis for pairwise comparison (pairs with P<0.05 are indicated). **(G)** ATAC-seq and histone ChIP-seq FE scores at pS22-Lamin A/C-binding sites, ordered by TSS distance from pS22-Lamin A/C-binding site center to closest transcription start site. pS22-Lamin A/C ChIP-seq signals are shown as a reference. See **Fig. S3A** for additional analysis on ATAC-seq and histone ChIP-seq. **(H)** Chromatin states of pS22-Lamin A/C-binding sites and Lamin A/C LADs. Chromatin states defined in normal dermal fibroblasts are used (Roadmap Epigenomics Consortium et al., 2015).

Phospho-mimetic Lamin C was far more enriched at pS22-Lamin A/C-binding sites than phospho-mimetic Lamin A (one-way ANOVA with post-hoc Tukey test P=1×10^-9^; **Fig. 2F; Fig. S2H**), consistent with the higher abundance of pS22-Lamin C compared with pS22-Lamin A in BJ-5ta (**Fig. 1D, E**). Together, these experiments revealed that S22/S392-phosphorylated Lamin A/C, most prominent in the form of S22/S392-phosphorylated Lamin C, has strong affinity to genomic sites outside of LADs.

### Phospho-S22-Lamin A/C physically associates with putative active enhancers

pS22-Lamin A/C-binding sites in the nuclear interior exhibited genomic features of active enhancers. First, a vast majority of the 22,966 pS22-Lamin A/C-binding sites in BJ-5ta fibroblasts were located distal to the transcription start sites (TSSs) of genes (89% outside of –1000 kb to +500 bp of TSSs) (**Fig. 2G**). Second, we performed ATAC-seq in BJ-5ta and observed that 88% of pS22-Lamin A/C-binding sites coincided with regions of accessible chromatin, a feature of regulatory regions (**Fig. 2B, G; Fig. S3A; Table S7**). Third, we performed H3K27ac ChIP-seq in BJ-5ta and observed that 82% of pS22-Lamin A/C-binding sites coincided with regions enriched for H3K27ac, a histone modification associated with active enhancers and promoters (**Fig. 2B, G; Fig. S3A; Table S8**). Fourth, we performed H3K4me3 ChIP-seq in BJ-5ta and observed that only 13% of pS22-Lamin A/C-binding sites coincided with regions enriched for H3K4me3, a histone modification associated with active promoters (**Fig. 2B, G; Fig. S3A; Table S9**). Together, the strong association of pS22-Lamin A/C-binding sites with H3K27ac but not H3K4me3 suggested a binding preference of pS22-Lamin A/C for enhancers over promoters. Fifth, comparison of the 22,966 pS22-Lamin A/C-binding sites with the chromatin state annotations in dermal fibroblasts (Roadmap Epigenomics Consortium et al., 2015) found that 59% of pS22-Lamin A/C-binding sites were located in regions annotated as “Enhancers” (empirical P<0.001 based on 2,000 permutations), whereas a smaller fraction of 16% were in “Active TSSs” (P<0.001) (**Fig. 2H**). Finally, although nearly all pS22-Lamin A/C-binding sites corresponded to accessible chromatin and H3K27ac-marked sites, pS22-Lamin A/C-binding sites corresponded to a minor subset of all accessible chromatin sites (26%) or all H3K27ac-marked sites (14%) in BJ-5ta (**Fig. S3A, B**). Thus, pS22-Lamin A/C physically associates with a specific subset of promoter-distal, enhancer-like elements.

### Phospho-S22-Lamin A/C-binding sites are strongly bound by the c-Jun transcription factor

Localization of pS22-Lamin A/C-binding sites at locations indicative of regulatory elements led us to investigate whether they are co-occupied by specific transcription factors (TFs). We performed *de novo* motif analysis and found that TF binding motifs for AP-1 (MEME E-value (Machanick and Bailey, 2011) E=1×10^-196^), FOX (E=6×10^-127^), and RUNX (E=2×10^-25^) were overrepresented within pS22-Lamin A/C-binding sites (+/–75 bp of the pS22-Lamin A/C-binding-site center), relative to all accessible chromatin sites identified by ATAC-seq in BJ-5ta (**Fig. 3A**). Among those motifs, the AP-1 motif was present at the highest frequency at pS22-Lamin A/C-binding sites (9,250 pS22-Lamin A/C-binding sites, 41%), with a peak frequency located at the center of pS22-Lamin A/C-binding sites (**Fig. 3A**). To test whether AP-1 transcription factors bind pS22-Lamin A/C-binding sites, we performed ChIP-seq in BJ-5ta for c-Jun, a core protein of the AP-1 dimeric transcription factors. c-Jun was strongly enriched at almost all pS22-Lamin A/C-binding sites (92% of 500-bp windows that contained pS22-Lamin A/C-binding sites; Fisher’s exact test P<2×10^-16^) (**Fig. 3B–D; Table S10**). Furthermore, c-Jun binding at pS22-Lamin A/C-binding sites was much stronger than that outside of pS22-Lamin A/C-binding sites (Mann-Whitney *U* test, P<2×10^-16^; **Fig. 3E**). We assessed the possibility that the strong co-association between pS22-Lamin A/C binding and c-Jun binding might be due to the high local chromatin accessibility of these co-associated sites. We stratified all 73,933 accessible sites defined by ATAC-seq in BJ-5ta into deciles by accessibility (i.e. ATAC-seq enrichment), and from each decile, randomly selected 100 pS22-Lamin A/C-*bound* accessible sites and 100 pS22-Lamin A/C-*unbound* accessible sites. In every accessibility decile, we observed more frequent c-Jun binding by ChIP-seq (Fisher’s exact test, P < 7×10^-4^; **Fig. S3C**) and stronger c-Jun enrichment levels (Mann-Whitney *U* test P=4×10^-13^ to 2×10^-23^; **Fig. 3F**) at pS22-Lamin A/C-bound accessible sites, compared with unbound sites. An analogous analysis with all 87,988 c-Jun-bound sites stratified into deciles by accessibility showed that, in every decile, c-Jun enrichment levels at pS22-Lamin A/C-*bound* sites was stronger than at pS22-Lamin A/C-*unbound* sites (Mann-Whitney *U* test P=1×10^-9^ to 5×10^-20^) (**Fig. S3D**). H3K27ac levels were also higher at accessible sites or c-Jun-binding sites with pS22-Lamin A/C binding versus without pS22-Lamin A/C binding, within the same accessibility decile (Mann-Whitney *U* test P < 6×10^-5^ among ATAC sites; P < 2×10^-14^ among c-Jun-binding sites; **Fig. 3F; Fig. S3D**). Corroborating these observations, pS22-Lamin A/C ChIP signals were strongly positively correlated with c-Jun ChIP signals (Pearson correlation coefficient *r*=0.63) at the 22,966 pS22-Lamin A/C-binding sites, which, for comparison, were stronger than the correlation with H3K27ac signals (*r*=0.31) or ATAC-seq signals (*r*=0.28) (**Fig. 3G**). The strong association between pS22-Lamin A/C binding and c-Jun binding suggests that c-Jun and pS22-Lamin A/C may function together at enhancer-like elements.

**Figure 3.**
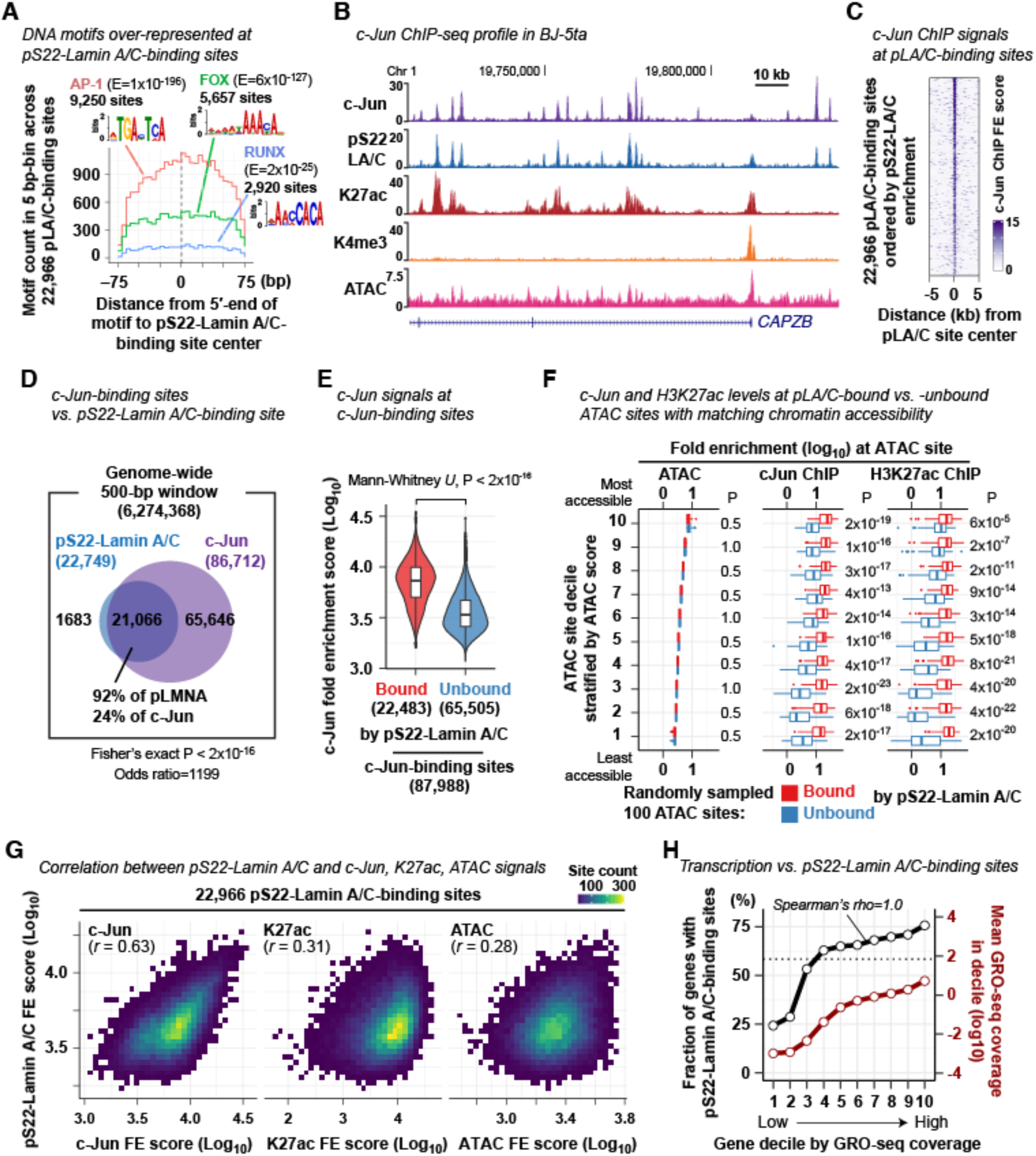
pS22-Lamin A/C association and c-Jun association are strongly correlated at pS22-Lamin A/C-binding sites. **(A)** DNA motif frequency at pS22-Lamin A/C-binding sites. E, motif occurrence probability score in *de novo* DNA motif search (Machanick and Bailey, 2011). **(B)** Representative c-Jun ChIP-seq fold-enrichment (FE) profile in BJ-5ta cells (performed in 3 biological replicates). Other ChIP- and ATAC-seq profiles are shown for comparison. **(C)** c-Jun ChIP-seq FE scores at pS22-Lamin A/C-binding sites. **(D)** Overlap between pS22-Lamin A/C-binding sites and c-Jun-binding sites. Number indicates the number of 500-bp windows that overlap pS22-Lamin A/C-binding sites and/or c-Jun-binding sites. **(E)** c-Jun ChIP FE scores at c-Jun-binding sites bound or unbound by pS22-Lamin A/C. Y-axis, sum of FE scores within +/–250 bp of site center. Box, interquartile range. Violin, kernel density. See **Methods** for box and violin plot notations used throughout the paper. **(F)** c-Jun ChIP and H3K27ac ChIP-seq FE scores at ATAC sites bound or unbound by pS22-Lamin A/C. ATAC sites (total 73,933 sites) are stratified into deciles by ATAC FE scores, and from each decile, 100 pS22-Lamin A/C-bound and 100 pS22-Lamin A/C-unbound ATAC sites are randomly selected, and their FE score distribution is visualized. Sum of FE scores within +/–250 bp of ATAC site center is used for analysis. Mann-Whitney *U*-test compares FE scores between pS22-Lamin A/C-bound and -unbound sites, and P-values are adjusted for multiple comparisons by the Benjamini-Hochberg method. See **Fig. S3C**, **D** for analyses on c-Jun-binding sites. **(G)** Two-dimensional histogram of cJun ChIP, H3K27ac ChIP, or ATAC-seq scores vs. pS22-Lamin A/C ChIP scores for the 22,966 pS22-Lamin A/C-binding sites. Sum of FE scores within +/–250 bp of site center is used. One square, one bin, with color grade representing the number of pS22-Lamin A/C-binding sites within the bin. *r*, pearson correlation coefficient. **(H)** (Black) Fraction of genes with pS22-Lamin A/C-binding sites (within gene body or 100 kb upstream) in gene decile stratified by GRO-seq coverage. Dotted line, fraction of all genes with pS22-Lamin A/C-binding sites. (Red) Mean GRO-seq coverage. GRO-seq was performed in BJ-5ta in 2 biological replicates. See **Fig. S3E, F** for analyses on genes further stratified by LAD overlap, gene lengths, and gene density.

### Phospho-S22-Lamin A/C-binding sites are located near highly-transcribed genes

We hypothesized that pS22-Lamin A/C-binding sites are located near genes undergoing active transcription given the enhancer features and c-Jun co-association of pS22-Lamin A/C-binding sites. We therefore performed GRO-seq to quantify transcriptional activity of genes in BJ-5ta cells and linked pS22-Lamin A/C-binding sites to gene transcription levels. We associated genes with pS22-Lamin A/C-binding sites that resided in the gene body or the 100-kb upstream region. The fraction of genes linked to pS22-Lamin A/C-binding sites was highest among the top 10% of highly transcribed genes (76%) and lowest among the bottom 10% of transcribed genes (24%), with strong positive monotonic relationship between the transcription levels and the fraction of genes with pS22-Lamin A/C-binding sites (Spearman’s rank correlation coefficient rho=1.0; **Fig. 3H**). The strong correlation remained in analysis of genes specifically located outside of LADs (rho=0.98; **Fig. S3E**), excluding the possibility that the observed positive correlation was driven by the localization of pS22-Lamin A/C binding outside of transcriptionally-inactive LADs. We further excluded the potential confounding effects of gene lengths and gene density (**Fig. S3F**). Together, these analyses revealed that pS22-Lamin A/C binding is associated with high transcriptional activity of local genes.

### Genes abnormally up-regulated in progeria-patient fibroblasts are relevant to progeria phenotypes

The heterozygous *LMNA* mutations that cause Hutchinson-Gilford progeria encode a mutant Lamin A protein, entitled progerin, which lacks part of the C-terminal domain that mediates chromatin and protein interactions (Eriksson et al., 2003; Bruston et al., 2010; Simon and Wilson, 2013; Gordon et al., 2014). Progerin interacts with wild-type Lamin A and Lamin C with high affinity and alters normal functions of wild-type Lamin A/C (Lee et al., 2016). The prevailing models for progerin’s pathological activity include disruption of Lamin A/C function at LADs (Gordon et al., 2014). We hypothesized that progerin may alter pS22-Lamin A/C function at enhancers. We therefore investigated the possibility that altered pS22-Lamin A/C binding to enhancers contributed to gene expression changes in progeria. We performed transcriptome analysis to identify genes dysregulated in fibroblast cell lines from progeria patients. We performed RNA-seq on primary dermal fibroblasts from two progeria patients (AG11498 and HGADFN167) and two normal individuals with similar ages (GM07492 and GM08398) (**Fig. S4A–C**). In addition, we obtained public RNA-seq data sets for primary dermal fibroblasts from ten progeria patients (AG11513, HGADFN188, HGADFN127, HGADFN164, HGADFN169, HGADFN178, HGADFN122, HGADFN143, HGADFN367, and HGADFN167) and ten normal individuals with similar ages (GM00969, GM05565, GM00498, GM05381, GM05400, GM05757, GM00409, GM00499, GM00038, and GM08398) (Fleischer et al., 2018) (**Fig. S4A**). By principal component analysis, RNA-seq datasets of the progeria patients were separated from those of the normal individuals along the second principal component, indicating a common gene expression signature among progeria-patient fibroblasts distinct from that of normal individuals (**Fig. 4A; Fig. S4D**; the first principal component corresponded to the study origin of the samples). Comparison between the 14 progeria RNA-seq data sets (for 11 distinct cell lines) and the 12 normal RNA-seq data sets (for 10 distinct cell lines after removing 2 outlier datasets; **Fig. S4D**) identified 1,117 dysregulated genes, 615 up-regulated in the progeria cell lines (“progeria-up” genes) and 502 down-regulated in the progeria cell lines (“progeria-down” genes) (**Fig. 4B, C; Table S11**). The progeria-up genes were strongly over-represented for specific DisGeNET-curated disease ontology terms (Piñero et al., 2017) (17 terms with P<0.001) that were well-documented progeria phenotypes, such as “infraction, middle cerebral artery” (Hypergeometric test (Zhou et al., 2019) P=7×10^-6^) (Silvera et al., 2013), “coronary artery disease” (P=2×10^-5^) (Olive et al., 2010), “cardiomegaly” (P=4×10^-5^) (Prakash et al., 2018), and “hypertensive disease” (P=1×10^-4^) (Merideth et al., 2008), and that were closely related to progeria phenotypes, such as “juvenile arthritis” (P=1×10^-5^) (Gordon et al., 2007) and “Congenital hypoplasia of femur” (P=1×10^-4^) (Gordon et al., 2007) (**Fig. 4D**). The progeria-down genes were associated with only three DisGeNet disease terms (P<0.001), including two terms that might be related to progeria phenotypes including “hip joint varus deformity” (Gordon et al., 2007) (P=6×10^-4^) and “short stature, mild” (P=9×10^-4^) (Gordon et al., 2011) (**Fig. 4D**). Thus, genes dysregulated in progeria-patient fibroblast lines were associated with the clinical components of progeria, with stronger association observed for genes upregulated in progeria fibroblasts.

**Figure 4.**
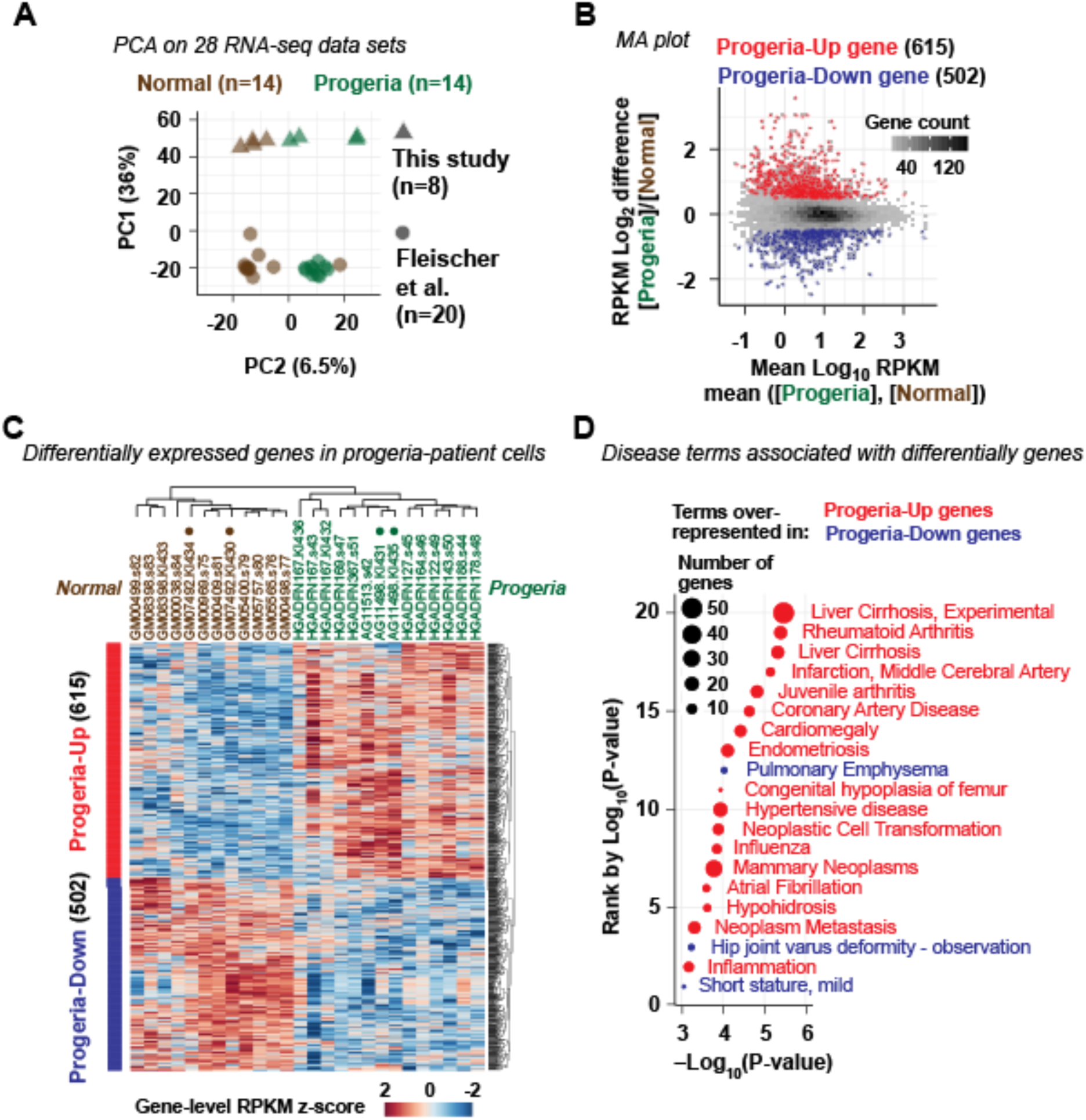
Genes abnormally up-regulated in progeria fibroblasts are relevant to progeria phenotypes. **(A)** Principal component analysis (PCA) on RNA-seq data sets of primary fibroblasts derived from normal individuals (14 data sets) and progeria patients (14 data sets). Percentage indicates the proportion of variance explained by the principal component. See **Fig. S4** for additional PCA analyses and sample details. **(B)** MA plot comparing mean RNA-seq RPKM scores across 14 progeria fibroblasts and those across 12 normal fibroblasts. **(C)** Hierarchical clustering of genes upregulated in progeria (“progeria-up”) and genes downregulated in progeria (“progeria-down”) (row) and normal and progeria fibroblast RNA-seq data sets (column). Solid circles, normal GM07492 and progeria AG11497 fibroblasts used in LAD and pS22-Lamin A/C analyses. **(D)** DisGeNet-curated disease terms overrepresented among progeria-up (red) and progeria-down genes (blue). Shown are terms with P < 0.001.

### LAD alterations do not explain most gene expression alterations in progeria

We next investigated the cause of the gene expression alterations in the progeria-patient fibroblasts. A prevailing model for progeria pathogenesis is that disruption of LADs causes loss of heterochromatin-associated histone modifications, which in turn alters gene expression (Shumaker et al., 2006; McCord et al., 2012; Gordon et al., 2014). To determine the extent to which LAD alterations in progeria fibroblasts could explain the observed gene expression changes, we performed ChIP-seq using anti-pan-N-terminal-Lamin A/C antibody in the progeria-patient fibroblast cell line AG11498 and the normal-individual fibroblast cell line GM07492 (**Fig. 5A**); these cell lines demonstrated transcriptomes representative of progeria versus normal fibroblasts, respectively (**Fig. 4C**). We identified 2,735 total or ‘union’ LADs, i.e., present in either the progeria or the normal fibroblast or both. Of those, 635 LADs (23%) were unique to either the normal or progeria fibroblasts: 353 LADs (13%) were present in normal fibroblasts but absent in progeria fibroblasts (called “lost LADs”), whereas 282 LADs (10%) were absent in normal fibroblasts but present in progeria fibroblasts (called “gained LADs”) (**Fig. 5A, B; Fig. S5A, B; Table S13**). To determine if gained or lost LADs were associated with changes of the heterochromatin-associated histone modifications H3K9me3 and H3K27me3, we performed ChIP-seq for H3K9me3 and H3K27me3 in the progeria cell line AG11498 and the normal cell line GM07492. H3K9me3 levels were reduced in the progeria cell line at lost LADs (Mann-Whitney *U* test P=3×10^-12^), but increased at gained LADs (P=3×10^-12^) (**Fig. 5C**), showing a positive correlation with the direction of LAD changes. H3K27me3 levels were reduced in progeria at both lost and gained LADs compared with the union LADs (P=1×10^-5^ for lost LADs and P=5×10^-7^ for gained LADs) (**Fig. 5D**), consistent with the previous report that H3K27me3 levels are globally reduced at gene-poor regions in progeria-patient fibroblasts (McCord et al., 2012). These observations confirm the previous findings that the LAD profile was altered in progeria and that LAD alterations in progeria are associated with changes of heterochromatin-associated histone modifications (Shumaker et al., 2006; McCord et al., 2012).

**Figure 5.**
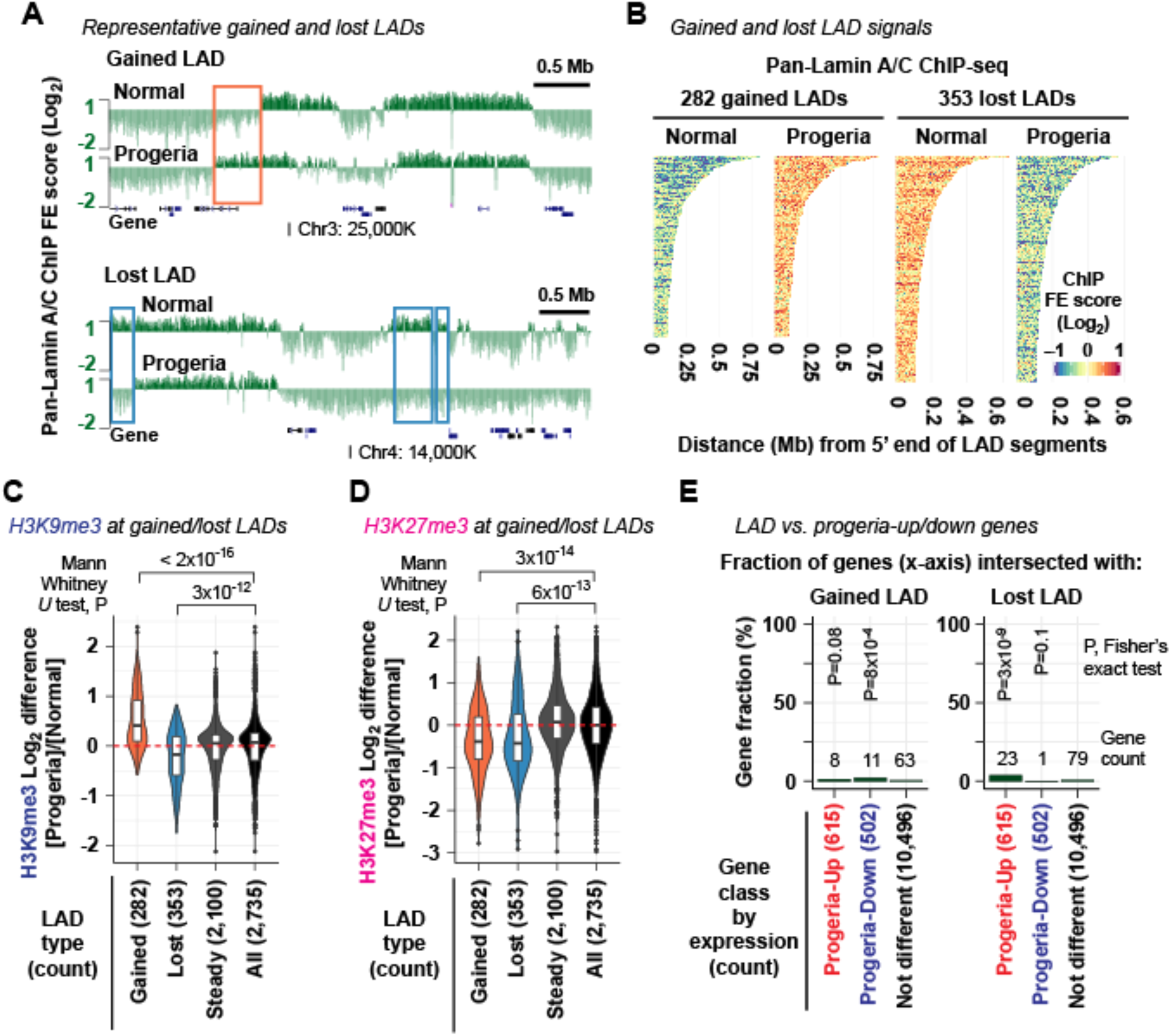
LAD alterations do not explain the majority of gene expression changes in progeria. **(A)** Representative pan-N-terminal-Lamin A/C ChIP-seq profiles in normal (GM07492) and progeria-patient (AG11498) fibroblasts (performed in 2 biological replicates). Log_2_ scale on the y axis. Square, representative gained LADs (present in progeria and absent in normal) and lost LADs (present in normal and absent in progeria). See **Fig. S5A, B** for additional gained and lost LADs. **(B)** Pan-N-terminal-Lamin A/C ChIP-seq FE score in the normal GM07492 and progeria AG11498 fibroblasts at gained and lost LADs in progeria. **(C)** H3K9me3 ChIP-seq FE log_2_ difference between normal (GM07492) and progeria (AG11498) fibroblasts (performed in 2 biological replicates). Red dashed line, Log_2_ difference equals 0 for reference. Mean FE scores in LADs are compared. **(D)** Same as **(C)**, but H3K27me3 log_2_ difference is shown. **(E)** Fraction of progeria-up and progeria-down genes whose gene body or upstream 100-kb region intersected with gained LADs (left) or lost LADs (right). P, Fisher’s exact test P value for association between being differentially expressed and being affiliated with LADs. See **Fig. S5C, D** for disease ontology analyses.

We examined the extent to which changes in LAD structure were associated with gene expression changes in progeria. We examined genes whose gene body or upstream 100-kb region intersected with lost or gained LADs. We asked if genes upregulated in progeria were affiliated with lost LADs or genes downregulated in progeria were affiliated with gained LADs. Only 23 out of 615 genes up-regulated in progeria intersected with lost LADs (**Fig. 5E**; **Table S14**). This subset accounted for only 3.7% of progeria-up genes (Fisher’s exact test P=3×10^-9^). Similarly, only 11 out of 502 genes downregulated in progeria intersected with gained LADs (**Fig. 5E**). This subset accounted for only 2.2% of progeria-down genes (P=8×10^-4^). Furthermore, these progeria-up genes in lost LADs (23 genes) or progeria-down genes in gained LADs (11 genes) were not over-represented for disease ontology terms linked to progeria phenotypes (**Fig. S5C, D**). These data revealed that local LAD alterations could not explain the vast majority of gene expression changes in progeria, either upregulation of gene expression by LAD losses or downregulation of gene expression by LAD gains.

### New pS22-Lamin A/C-binding sites emerge in normally quiescent loci in progeria-patient fibroblasts

We hypothesized that the gene expression changes observed in the progeria-patient fibroblasts were associated with alterations of pS22-Lamin A/C-binding sites, given that they were largely not associated with the disruption of LADs. We observed that pS22-Lamin A/C was present in the interior of interphase nuclei of the progeria-patient fibroblast cell line AG11498, but not at the nuclear periphery, as it is in the normal fibroblast cell line GM07492 (**Fig. S6A**). We found that the major isoform phosphorylated at S22 in the progeria-patient and normal fibroblast cells was Lamin C, while S22-phosphorylation of progerin was not detectable in Western blotting in the progeria-patient fibroblasts (**Fig. S6B**). We performed pS22-Lamin A/C ChIP-seq in the progeria-patient fibroblast cell line AG11498 and the normal-individual fibroblast cell line GM07492 (the same cell lines used in pan-N-terminal-Lamin A/C ChIP-seq). We identified a union set of 15,323 pS22-Lamin A/C-binding sites found in either the normal-individual fibroblast GM07492 or the progeria-patient fibroblast AG11498 or both. We observed significant alteration of the pS22-Lamin A/C-binding site profile in the progeria fibroblasts: of the 15,323 union pS22-Lamin A/C-binding sites, 2,796 pS22-Lamin A/C-binding sites (18%) were specific to the progeria fibroblast line (termed “gained pS22-Lamin A/C-binding sites”) whereas 2,425 pS22-Lamin A/C-binding sites (16%) were specific to the normal fibroblast line (termed “lost” pS22-Lamin A/C-binding sites) (**Fig. 6A, B; Fig. S6C, D; Table S12**). Gained pS22-Lamin A/C-binding sites were highly over-represented within the “quiescent” chromatin-state annotation derived from normal dermal fibroblasts (Roadmap Epigenomics Consortium et al., 2015) (25% of gained pS22-Lamin A/C-binding sites vs. 10% of all pS22-Lamin A/C-binding sites; Fisher’s exact test P=2×10^-144^), whereas lost pS22-Lamin A/C-binding sites were over-represented in “enhancer” regions (72% of lost vs. 62% of all pS22-Lamin A/C-binding sites; P=8×10^-30^) (**Fig. 6C**). Thus, new pS22-Lamin A/C-binding sites emerged in locations in progeria, a subset of which possessed chromatin features of quiescence in normal fibroblasts, while a subset of the wild-type enhancer pS22-Lamin A/C-binding sites were lost in progeria. The absence of detectable pS22-progerin (**Fig. S6B**) suggests that progerin disrupted the binding specificity of wild-type pS22-Lamin A/C in progeria fibroblasts.

**Figure 6.**
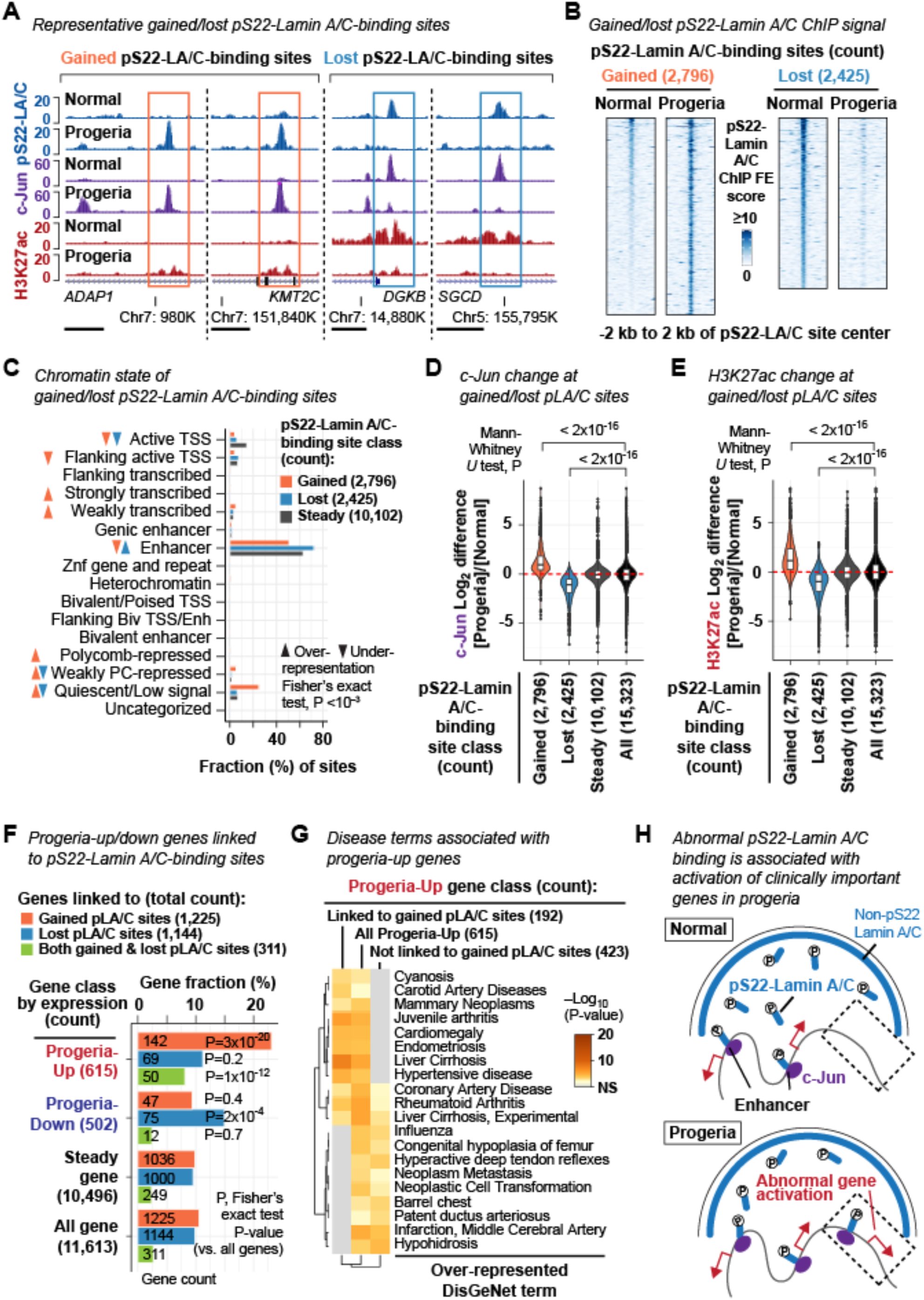
New pS22-Lamin A/C-binding sites emerged in progeria are associated with up-regulation of genes relevant to progeria phenotypes. **(A)** pS22-Lamin A/C ChIP-seq profiles highlighting representative gained and lost pLamin A/C-binding sites (performed in 2 biological replicates). Normal, GM07492 fibroblast. Progeria, AG11498 fibroblast. Horizontal bar, 2 kb. See **Fig. S6** for subcellular localization and abundance of pS22-Lamin A/C and additional characterization of pS22-Lamin A/C-binding sites in progeria fibroblasts. **(B)** pS22-Lamin A/C ChIP-seq FE score in the normal GM07492 and progeria AG11498 fibroblasts at gained and lost pS22-Lamin A/C-binding sites in progeria. **(C)** Chromatin state of gained and lost pS22-Lamin A/C-binding sites. Fisher’s exact test assesses association between being called as gained or lost and being affiliated with each state. **(D)** c-Jun ChIP-seq FE score difference between normal (GM07492) and progeria (AG11498) fibroblasts (performed in 2 biological replicates). Red dashed line, log_2_ difference equals 0 for reference. Sum of FE scores within +/–250 bp of site center is compared. **(E)** Same as **(D)**, but FE scores of H3K27ac ChIP-seq (performed in 2 biological replicates) are shown. **(F)** Fraction of progeria-up and progeria-down genes linked to gained and/or lost pS22-Lamin A/C-binding sites. Genes are linked to pS22-Lamin A/C-binding sites when pS22-Lamin A/C-binding sites reside in the gene body or the upstream 100-kb region. **(G)** DisGeNet-curated disease terms over-represented among progeria-up genes linked or not linked to gained pS22-Lamin A/C-binding sites. See **Fig. S6H** for progeria-down genes. **(H)** Summary. In normal fibroblasts, pS22-Lamin A/C associates with putative active enhancers. In progeria fibroblasts, gains of pS22-Lamin A/C-binding sites in abnormal locations accompany activation of genes relevant to progeria phenotypes.

We hypothesized that changes in pS22-Lamin A/C-binding sites between progeria and normal fibroblasts reflected changes in the activity level of active enhancers. Because the level of pS22-Lamin A/C binding was positively correlated with c-Jun and H3K27ac levels (**Fig. 3**), we hypothesized that gains of pS22-Lamin A/C-binding sites in progeria may accompany gains in c-Jun and H3K27ac levels, whereas losses of pS22-Lamin A/C-binding sites may accompany losses of c-Jun and H3K27ac levels. To test this hypothesis, we performed c-Jun and H3K27ac ChIP-seq in the progeria cell line AG11498 and normal cell line GM07492. At the 2,796 gained pS22-Lamin A/C-binding sites, c-Jun and H3K27ac levels were strongly elevated in the progeria cell line relative to the normal cell line, as compared with all 15,323 union pS22-Lamin A/C-binding sites (**Fig. 6D, E; Fig. S6E, F**; Mann-Whitney *U* test P<2×10^-16^ for both c-Jun and H3K27ac). In contrast, at the 2,425 lost pS22-Lamin A/C-binding sites, c-Jun and H3K27ac levels were strongly diminished in the progeria cell line relative to the normal cell line (**Fig. 6D, E; Fig. S6E, F**; P<2×10^-16^ for both c-Jun and H3K27ac). Thus, the alteration of the pS22-Lamin A/C binding profile in progeria fibroblasts was accompanied by local alterations of c-Jun binding and H3K27ac levels at the pS22-Lamin A/C-binding sites, suggesting a link between the alteration of the pS22-Lamin A/C binding profile and transcriptional dysregulation in progeria fibroblasts.

### Gains of pS22-Lamin A/C binding in progeria accompany abnormal transcriptional activation of genes clinically important to progeria pathophysiology

We hypothesized that the gains and losses of pS22-Lamin A/C-binding sites in progeria, and the associated changes in c-Jun and H3K27ac levels at those sites, affected transcription in progeria fibroblasts. To test this, we linked pS22-Lamin A/C-binding sites to genes, associating pS22-Lamin A/C-binding sites that reside in the gene body or the 100-kb upstream region of candidate target genes. By this metric, 85% of the 15,323 union pS22-Lamin A/C-binding sites were linked to at least one gene (total 6229-linked genes out of the 11,613 all genes). Of those, 1,225 genes were linked only to pS22-Lamin A/C-binding sites gained in progeria (11% of all genes), and 1,144 genes were linked only to pS22-Lamin A/C-binding sites lost in progeria (9.9% of all genes) (**Fig. S6G; Table S11**). Genes linked only to gained pS22-Lamin A/C-binding sites were highly over-represented among genes up-regulated in progeria (142 genes, 23%, P=3×10^-20^), but not among genes down-regulated in progeria (47 genes, 9.3%, P=0.4) (**Fig. 6F**). Genes linked only to lost pS22-Lamin A/C-binding sites were over-represented among genes down-regulated in progeria (75 genes, 15%, P=2×10^-4^), but not among genes upregulated in progeria (69 genes, 11%, P=0.2) (**Fig. 6F**). Thus, in progeria-patient fibroblasts, gains of pS22-Lamin A/C binding were associated with up-regulation of genes, and losses of pS22-Lamin A/C binding were associated with down-regulation of genes. Interestingly, 311 genes were linked to both gained and lost pS22-Lamin A/C-binding sites (2.7% of all genes), but were only over-represented among genes upregulated in progeria (50 genes, 8.1%, P=1×10^-12^), but not among genes downregulated in progeria (12 genes, 2.3%, P=0.7) (**Fig. 6F**), suggesting a dominant association between gains of pS22-Lamin A/C-binding and up-regulation of genes in progeria.

We hypothesized that gained or lost pS22-Lamin A/C-binding sites at enhancers linked to genes dysregulated in progeria may be relevant to progeria phenotypes. The progeria-up genes linked to gained pS22-Lamin A/C-binding sites (192 genes) were highly over-represented for DisGeNet disease ontology terms relevant to progeria phenotypes such as carotid artery disease (Hypergeometric test (Zhou et al., 2019) P=0.007) (Gerhard-Herman et al., 2012), juvenile arthritis (P=1×10^-5^) (Gordon et al., 2007), hypertensive disease (P=0.0001) (Merideth et al., 2008), and cardiomegaly (P=4×10^-5^) (Prakash et al., 2018) (**Fig. 6G**). None of these terms were overrepresented among the progeria-up genes not linked to a pS22-Lamin A/C-binding site (423 genes) (**Fig. 6G**). Consistent with the ontology analysis, progeria-up genes with gained pS22-Lamin A/C binding included important cardiovascular genes (**Table S14**). Examples include *FHL1*, overexpression of which causes myopathies (Schessl et al., 2008) and mutations of which causes Emery-Dreifuss muscular dystrophy also caused by *LMNA* mutations (Windpassinger et al., 2008; Gueneau et al., 2009) (**Fig. S6D**). Unlike progeria-up genes linked to gained pS22-Lamin A/C-binding sites, the progeria-down genes linked to lost pS22-Lamin A/C-binding sites were not associated with disease ontology terms relevant to progeria phenotypes (**Fig. S6H**). Thus, gained pS22-Lamin A/C-binding sites mark a subset of abnormally activated genes in progeria that are highly relevant to progeria phenotypes. We propose that misdirection of pS22-Lamin A/C to normally quiescent genomic locations is a novel mechanism for the transcriptional activation of pathogenesis-related gene pathways in progeria (**Fig. 6H**).

## DISCUSSION

Nuclear lamins have been extensively studied in the context of the nuclear lamina (Dechat et al., 2010b; van Steensel and Belmont, 2017). In this paper, we investigated the genomic localization and function of Ser22-phosphorylated nuclear lamin A/C (pS22-Lamin A/C) localized to the interior of the nucleus. pS22-Lamin A/C associated with kilobase-wide sites characteristic of active enhancers, in stark contrast to nuclear-peripheral Lamin A/C, which associates with megabase-wide heterochromatin domains (Meuleman et al., 2012; Lund et al., 2014). The existence of Lamin A/C in the nuclear interior has been documented for decades (Goldman et al., 1992; Bridger et al., 1993; Moir et al., 1994; Hozák et al., 1995; Barboro et al., 2002; Naetar et al., 2008; Shimi et al., 2008; Swift et al., 2013; Buxboim et al., 2014; Kochin et al., 2014; Gesson et al., 2016), yet the specific function of nuclear-interior Lamin A/C had been elusive. This report demonstrates that pS22-Lamin A/C-binding sites were affiliated with the signature of enhancers, were frequently co-occupied by the AP-1 transcription factor c-Jun, a strong transcriptional activator, and were positively correlated with transcriptional activity of local genes. Furthermore, gains and losses of pS22-Lamin A/C-binding sites in progeria-patient cells were accompanied by increased or decreased local H3K27ac and up-or down-regulation of nearby genes, respectively. These observations provide evidence that nuclear-interior pS22-Lamin A/C functions as a positive modulator of enhancer activity and that the enhancer function of pS22-Lamin A/C may be altered in progeria and contribute to progeria pathogenesis.

Decades of work have established that nuclear lamins at the nuclear lamina are associated with transcriptionally-inactive regions (Pickersgill et al., 2006; Guelen et al., 2008; Ikegami et al., 2010; Meuleman et al., 2012; van Steensel and Belmont, 2017). In this respect, the data showing that pS22-Lamin A/C-binding sites exhibit the characteristics of active enhancers and are associated with active transcription is striking. Our observation that pS22-Lamin C was more abundant than pS22-Lamin A and that phospho-mimetic Lamin C, but not phospho-mimetic Lamin A, strongly bound pS22-Lamin A/C-binding sites suggest that enhancer binding may be a distinctive feature of Lamin C as compared with Lamin A. While specific functions and localization of Lamin C are unknown, Lamin C lacks the C-terminal sequence present in Lamin A that undergoes farnesylation, a modification thought to promote nuclear lamina incorporation of Lamin A (Dechat et al., 2008). Thus, the lack of farnesylation in combination with S22/S392 phosphorylation may promote a nucleoplasmic Lamin C fraction that binds to enhancer-like elements. Our finding that chromatin accessibility is not a strong predictor of the location of pS22-Lamin A/C-binding sites suggests that distinct targeting mechanisms exist to direct pS22-Lamin A/C to putative active enhancers. We observed that pS22-Lamin A/C enrichment is strongly correlated with the enrichment of the AP-1 transcription factor c-Jun. Lamin A/C is known to interact with c-Fos, the binding partner of c-Jun in the functional AP-1 complex (Ivorra et al., 2006; González et al., 2008). The AP-1 transcription factor complex is therefore a candidate for functioning to target pS22-Lamin A/C to specific putative enhancers.

Recently, nuclear-interior Lamin A/C has been studied in the context of its interaction with lamina-associated polypeptide 2 alpha (LAP2ɑ) (Naetar et al., 2008; Dechat et al., 2010a). One report suggested that LAP2ɑ-interacting Lamin A/C associates with megabase-wide genomic regions that lie at euchromatin regions without specific localization at promoters or enhancers (Gesson et al., 2016). pS22-Lamin A/C binds to specific enhancer-like elements and promoters, and therefore, is likely distinct from LAP2ɑ-interacting Lamin A/C.

Altered gene expression programs have been reported in fibroblasts and other cell types derived from progeria patients (Csoka et al., 2004; Zhang et al., 2011), yet the underlying mechanisms have remained unclear. A prevailing hypothesis suggested that progerin, the mutant Lamin A/C protein expressed in progeria, accumulates at the nuclear lamina due to its permanent farnesylation and disrupts normal interactions between Lamin A/C and LADs, causing heterochromatin disorganization, and, in turn, altered expression of genes located in LADs (Shumaker et al., 2006; McCord et al., 2012). One limitation of this model is that it does not explain the specific and abundant gene expression changes that occur outside of LADs. We propose a novel hypothesis, in which mis-direction of pS22-Lamin A/C to otherwise unbound enhancer regions or quiescent regions results in abnormal direct transcriptional activation of genes relevant to progeria pathogenesis. Because progerin itself did not appear to be phosphorylated at S22, the direct interaction between progerin and Lamin A/C (Lee et al., 2016) may contribute to mis-direction of pS22-Lamin A/C in progeria. Understanding the molecular mechanisms for how pS22-Lamin A/C is mis-localized in progeria-patient cells is an important area for future investigations.

pS22-Lamin A/C has been regarded as a byproduct of mitotic nuclear envelope breakdown (Gerace and Blobel, 1980) or as a pool of disassembled lamins to be degraded when the nuclear lamina is compromised by mechanical stress (Swift et al., 2013; Buxboim et al., 2014). Our data that pS22-Lamin A/C is present throughout the cell cycle during unperturbed cellular conditions, localized at a specific subset of putative active enhancers, and associated with transcriptional alterations in progeria, suggested that pS22-Lamin A/C is a previously-unrecognized functional species of Lamin A/C in the interior of the nucleus. Much like the function of nuclear lamins in transcriptional repression at the nuclear lamina remains under active investigation (Leemans et al., 2019), the causal relationship between pS22-Lamin A/C binding at putative enhancers and transcriptional regulation remains to be determined. Regardless, the characteristics of pS22-Lamin A/C unveiled in this study offer a new foundation for investigating the functions of Lamin A/C, its role in transcriptional regulation, and the mechanisms underlying human degenerative disorders caused by *LMNA* mutations.

## ACKNOWLEDGEMENTS

We thank technical assistance from the University of Chicago Functional Genomics Core, the University of Chicago Cytometry and Antibody Technology Core, the University of Chicago Light Microscopy Core, and Princeton University Genomics Core Facility. We acknowledge Life Science Editors for editorial assistance. This work was funded by NIH R21/R33 AG054770 (K.I. and I.P.M.), NIH R01 HL147571 and R01 HL148719 (I.P.M.), and the Progeria Research Foundation grant #2009-0028 (K.I. and J.D.L.).

## AUTHOR CONTRIBUTIONS

K.I., J.D.L., I.P.M. conceived the study. K.I., S.S., and O.A. performed the experiments. K.I. analyzed the data and wrote the manuscript. J.D.L. and I.P.M. provided intellectual contributions and participated in manuscript writing.

## STAR METHODS

### LEAD CONTACT AND MATERIALS AVAILABILITY

Further information and requests for resources and reagents should be directed to and will be fulfilled by the Lead Contact, Kohta Ikegami.

### EXPERIMENTAL MODEL AND SUBJECT DETAILS

BJ-5ta (ATCC catalog # CRL-4001) is a TERT-immorta reskin of male neonate) that retains normal fibroblast cell growth phenotypes and does not exhibit transformed phenotypes (Jiang et al., 1999). Generation of the BJ-5ta-derived *LMNA^−/−^*cell line and *LMNA^−/−^* cells expressing wild-type and mutant *LMNA* transgenes is described in the following sections. Primary dermal fibroblasts used are GM07492 (source: thigh of 17-year-old male normal individual, Coriell Cell Repository), GM08398 (source: inguinal area of 8-year-old male normal individual, Coriell Cell Repository), AG11498 (source: thigh of 14-year-old male Hutchinson-Gilford progeria patient, Coriell Cell Repository), and HGADFN167 (source: posterior lower trunk of 8-year-old male Hutchinson-Gilford progeria patient, Progeria Research Foundation) (**Table S2**). We verified that, at the time of cell harvest, the progeria and control cells were not undergoing senescence (beta-galactosidase positive cells < 5%), a feature that late-passage progeria cells could manifest (Sieprath et al., 2015) (**Fig. S4B, C**). All cells were cultured in standard cell-culture-treated plastic dishes unless otherwise noted. BJ-5ta and its derivatives were cultured in high-glucose DMEM (Gibco, 11965-092) containing 9% fetal bovine serum (FBS), 90 U/mL penicillin, 90 µg/mL streptomycin streptomycin at 37°C under 5% CO_2_. Primary skin fibroblasts were cultured in MEM Alpha (Gibco, 12561-056) containing 9% fetal bovine serum (FBS), 90 U/mL penicillin, 90 µg/mL streptomycin streptomycin at 37°C under 5% CO_2_.

## METHOD DETAILS

### Cell synchronization by thymidine block

BJ-5ta cells in the DMEM growth medium were maintained at confluency for 2 days (G0 arrest by contact inhibition) and then passaged to a culture plate at a low density in the DMEM growth medium supplemented with 2 mM thymidine (Sigma-Aldrich T9250) for 17 hours. This allowed cells to re-enter into the G1 stage of the cell cycle and become arrested at the G1/S boundary. Cells were then washed and cultured with the growth medium with 2.5 µM deoxycytidine (without thymidine) and harvested at 0 (i.e. G1/S-arrested cells), 4, 6, 8, 10, 12, and 14 hours later. As a reference, G0-arrested cells were released in the growth medium without thymidine and harvested 14 hours later (“asynchronous” cells).

### Generation of LMNA^−/−^ cell line

We cloned DNA sequences for sgRNA1 (oligonucleotides KI223 and KI224; **Table S1**; PAM, forward strand at chr1:156,084,863-156,084,866 in hg19) or sgRNA3 (oligonucleotides KI227 and KI228, **Table S1**; PAM, forward strand at chr1:156,084,953-156,084,956 in hg19), targeting the exon 1 of *LMNA*, into the all-in-one lentivirus vector LentiCRISPRv2 (a gift from Feng Zhang; Addgene plasmid # 52961) (Sanjana et al., 2014). The cloned lentiCRISPR vectors were individually transfected to HEK293FT cells with the packaging vectors psPAX2 (gift from Didier Trono; Addgene plasmid #12260) and pCMV-VSV-G (a gift from Bob Weinberg; Addgene plasmid #8454) (Stewart et al., 2003) to produce lentivirus. A mixture of the lentiviral tissue-culture supernatant for sgRNA1 and sgRNA3 (each at 0.25 dilution) was applied to BJ-5ta cells in the presence of 7.5 µg/mL polybrene for transduction. Successfully transduced cells were selected by 3 µg/mL puromycin and seeded to 10-cm dishes with a density of 100 cells per dish. Clonal populations were expanded and analyzed by western blotting for Lamin A and Lamin C protein expression. The clone cc1170-1AD2 lacks Lamin A and Lamin C protein expression, has nullizygous frameshift mutations, and is used in this study.

### Wild-type and phospho-mutant Lamin A/C expression in LMNA^−/−^ cells

We cloned Lamin A or Lamin C cDNAs with S22 and S392 mutations or without mutations into the all-in-one doxycycline inducible lentivirus vector pCW57-MCS1-P2A-MCS2-PGK-Blast (gift from Adam Karpf; Addgene plasmid #80921) (Barger et al., 2019) using HiFi assembly (New England Biolabs E2621). DNA fragments containing S22D (TCG to GAC) and S22A (TCG to GCT) mutations were chemically synthesized (Integrated DNA Technologies, Inc). DNA fragments containing S392D (AGC to GAC) and S392A (AGC to GCC) mutations were amplified by PCR using published Lamin A/C cDNA plasmids containing these mutations as a template (Kochin et al., 2014) (gifts from Drs. Robert Goldman and John E Eriksson). The Lamin A/C expression vectors were designed to harbor silent mutations at the sgRNA1 and sgRNA3 CRISPR targeting sites (see *Generation of LMNA^−/−^ cell line*) to prevent the transgenes from being targeted by Cas9 in *LMNA^−/−^* cells. Lentivirus was produced as described above and transduced to BJ-5ta-derived *LMNA^−/−^*cells. The transduced cells were selected under 10 µg/mL blasticidin. The cell population IDs are cc1499-1 (wild-type Lamin A); cc1499-2 (Lamin A with S22A/S392A); cc1499-3 (Lamin A with S22D/S392D); cc1499-4 (wild-type Lamin C); cc1499-5 (Lamin C with S22A/S392A); and cc1499-6 (Lamin C with S22D/S392D). The transgene expression was induced by 20 µg/mL doxycycline for 72 hours. This doxycycline concentration did not affect overall growth of the generated cell lines.

### ELISA

Lamin A/C N-terminal aa2-30 peptides (ETPSQRRATRSGAQASSTPLSPTRITRLQ) with phosphorylated Ser22 (Lot U2312EI090-3/PE2186; purity 90.0%; MW 3234.45) or non-phosphorylated Ser22 (Lot U2312EI090-1/PE2183; purity 96.1%; MW 3154.47) were synthesized by Genscript (New Jersey, USA), and the quality was verified by the manufacturer. Peptides were immobilized to maleic anhydride-activated plastic wells (Pierce catalog 15100). Coated wells were blocked with 5% non-fat milk and 0.1% Tween 20 and then incubated with anti-phospho-Ser22-Lamin A/C antibody (Cell Signaling 13448S, Lot 1, 1:1000 dilution) or the anti-pan-N-terminal-Lamin A/C antibody (Santa Cruz Biotechnology sc-376248, Lot H2812, 1:5000) for 1 hour at 37°C. After washing, wells were incubated with horseradish peroxidase (HRP)-conjugated anti-rabbit IgG (GE Healthcare NA934V, Lot 9636020) or HRP-conjugated anti-mouse IgG (GE Healthcare NA931V, Lot 9648752). HRP activity was detected by 3,3′,5,5′-tetramethylbenzidine (TMB)-based colorimetric reaction (Pierce catalog #34022). The reaction was treated with 3N HCl, and the absorbance at 450 nm (reaction) and 550 nm (reference) was measured by a microplate reader. See **Quantification and Statistical Analysis** for downstream analyses.

### Western blot

Protein extract was separated by 4-12% Bis-Tris SDS-PAGE with MOPS buffer and transferred to a PVDF membrane. After blocking, the membranes were incubated for 12 hours or longer at 4°C with rabbit monoclonal anti-phospho-Ser22-Lamin A/C antibody D2B2E (Cell Signaling 13448S, Lot 1; 1:1000 dilution) or mouse monoclonal anti-pan-N-terminal-Lamin A/C antibody E1 (Santa Cruz Biotechnology sc-376248, Lot H2812; 1:1000 dilution). Primary antibodies were detected by HRP-conjugated anti-rabbit IgG (GE Healthcare NA934V, Lot 9636020; 1:10000 dilution) or HRP-conjugated anti-mouse IgG (GE Healthcare NA931V, Lot 9648752; 1:10000 dilution). Signals were produced by enhanced chemiluminescence (ECL) and detected digitally by a Bio-Rad ChemiDoc imager. The gel after protein transfer was counter-stained by coomassie to evaluate the loaded protein amount. See **Quantification and Statistical Analysis** for downstream analyses.

### Flow cytometry

For cell-cycle analysis by DAPI, detached cells were incubated with 70% cold ethanol for 12 hours or longer at –20°C for fixation. The fixed cells were incubated with FACS buffer (2% FBS, 1 mM EDTA, and 0.1% Tween 20 in PBS) for 10 min at room temperature. Cells were then incubated with PBS supplemented with 0.1% Triton and 1 µg/mL DAPI for 10 min at room temperature. The DAPI-stained cells were resuspended in FACS buffer and then analyzed by Fortessa 4-15 HTS or Fortessa X20 5-18 flow cytometry analyzers (BD Biosciences).

To stain cells for pS22-Lamin A/C, cells were fixed in PHEM buffer (60 mM PIPES-KOH pH7.5, 25 mM HEPES-KOH pH7.5, 10 mM EGTA, 4 mM MgSO_4_) supplemented with 4% formaldehyde, 0.5% Triton, and 100 nM phosphatase inhibitor Nodularin (Enzo ALX-350-061) for 15 min at 37°C. Cells were blocked with Blocking buffer (FACS buffer supplemented with 5% normal goat serum) and incubated with Alexa-647-conjugated rabbit monoclonal anti-phospho-Ser22-Lamin A/C antibody D2B2E (labeled at Cell Signaling, product ID 97262BC, Lot 1, 1:30 dilution) in Blocking buffer for 1 hour at 37°C. Cells were counter-stained with 1 µg/mL DAPI in FACS buffer. The stained cells were analyzed by Fortessa 4-15 HTS or Fortessa X20 5-18 flow cytometry analyzers. See **Quantification and Statistical Analysis** for downstream analyses.

### Immunofluorescence

Cells were grown on uncoated glass coverslips under the standard culture condition (see *Cell Culture*). Cells were fixed in PHEM buffer (60 mM PIPES-KOH pH7.5, 25 mM HEPES-KOH pH7.5, 10 mM EGTA, 4 mM MgSO_4_) supplemented with 4% formaldehyde, 0.5% Triton, and 100 nM phosphatase inhibitor Nodularin (Enzo ALX-350-061) for 10 min at 37°C. Cells on coverslips were blocked with Blocking buffer (1% skim milk and 5% goat serum in PBS), and then incubated with primary antibodies in Blocking buffer. Antibodies used in immunofluorescence are: Alexa 647-conjugated rabbit monoclonal anti-phospho-Ser22-Lamin A/C antibody D2B2E (labeled at Cell Signaling, product ID 97262BC, Lot 1, 1:100 dilution), mouse monoclonal anti-pan-N-terminal-Lamin A/C antibody E1 (Santa Cruz Biotechnology sc-376248, Lot # H2812, 1:5000), or mouse monoclonal anti-full-length-Lamin A/C antibody 4C4 (Abcam ab190380, Lot GR201137-1; 1:1000). Cells were incubated with secondary antibodies, counterstained by DAPI, and cured in ProLong Gold mounting medium (Molecular Probes, P36930). Cells were imaged using a Leica SP8 confocal microscope with a 63x or 100x objective. See **Quantification and Statistical Analysis** for downstream analyses.

### Senescence-associated beta-galactosidase assay

Cells were grown on coverslips and fixed for 5 min in 2% formaldehyde and 0.2% glutaraldehyde in PBS. Cells were washed with PBS and incubated in X-gal staining solution (40 mM citric acid/sodium phosphate buffer, 5 mM K_4_[Fe(CN)_6_] 3H_2_O, 5 mM K_3_[Fe(CN)_6_], 150 mM NaCl, 2 mM MgCl_2_, and 1 mg/mL X-gal) for 16 h at 37°C. The coverslips were washed with PBS and mounted for microscopy. To determine the staining percentages, cells from 10 randomly selected areas at 20x magnification were counted.

### ChIP-seq

Cells in culture dishes were crosslinked in 1% formaldehyde for 15 min at room temperature, and the reaction was quenched by 125 mM glycine. Cross-linked cells were washed with LB1 (50 mM HEPES-KOH pH7.5, 140 mM NaCl, 1 mM EDTA, 10% glycerol, 0.5% NP40, 0.25% Triton X-100) and LB2 (200 mM NaCl, 1 mM EDTA, 0.5 mM EGTA, and 10 mM Tris-HCl pH 8.0). Cells were resuspended in LB3-Triton (1 mM EDTA, 0.5 mM EGTA, 10 mM Tris-HCl pH 8, 100 mM NaCl, 0.1% Na-Deoxycholate, 0.5% N-lauroyl sarcosine, 1% Triton) supplemented with 1x protease inhibitor cocktail (Calbiochem 539131) and 100 nM phosphatase inhibitor Nodularin (Enzo ALX-350-061), and chromatin was extracted by sonication. The cell extract was cleared by 14,000 g centrifugation for 10 min. An aliquot of cell extract was saved for input DNA sequencing. Cell extract from one million cells was incubated with antibodies in a 200-µL reaction for 12 hours or longer at 4°C. Antibodies used in ChIP are: rabbit monoclonal anti-phospho-Ser22-Lamin A/C antibody D2B2E (Cell Signaling 13448S, Lot 1; 5 µL per IP); mouse monoclonal anti-pan-N-terminal-Lamin A/C antibody E1 (Santa Cruz Biotechnology sc-376248, Lot H2812; 10 µL per IP); mouse monoclonal anti-full-length-Lamin A/C antibody 4C4 (Abcam ab190380, Lot GR201137-1; 4 µL per IP); rabbit polyclonal anti-c-Jun antibody (Santa Cruz Biotechnology sc-1694, Lot D1014; 20 µL per IP); mouse monoclonal anti-H3K27ac antibody (Wako MABI0309, Lot 14007; 2 µL per IP); mouse monoclonal anti-H3K4me3 antibody (Wako MABI14004, Lot 14004; 2 µL per IP); rabbit polyclonal anti-H3K9me3 antibody (Abcam ab8898, Lot GR232099-3; 2 µL per IP); and mouse monoclonal anti-H3K27me3 antibody (Active Motif MABI0323, Lot 17019020; 2 µL per IP). Immunocomplex was captured by Protein A-conjugated sepharose beads (for rabbit antibodies) or Protein G-conjugated magnetic beads (for mouse antibodies) and washed. Immunoprecipitated DNA was reverse-crosslinked and used to construct high-throughput sequencing libraries using NEBNext Ultra DNA Library Prep Kit (New England Biolabs, E7370). DNA libraries were processed on a Illumina HiSeq machine for single-end sequencing. See **Quantification and Statistical Analysis** for downstream analyses. ChIP-seq experiments are listed in **Table S3**.

### ATAC-seq

One hundred thousand trypsinized cells were incubated with ATAC hypotonic buffer (10 mM Tris pH 7.5, 10 mM NaCl, 3 mM MgCl_2_) for 10 min at 4°C during 500 g centrifugation. Cells were incubated in Tagmentation mix (Tagmentation DNA buffer Illumina 15027866; Tagmentation DNA enzyme Illumina 15027865) for 30 min at 37°C. Purified DNA was used to construct high-throughput sequencing libraries using NEBNext High-Fidelity 2x PCR Master Mix (New England Biolabs M0541). DNA libraries were processed on a Illumina NextSeq machine for paired-end 41-nt sequencing. See **Quantification and Statistical Analysis** for downstream analyses. ATAC-seq experiments are listed in **Table S3**.

### RNA-seq

Total RNAs were purified by Trizol LS (Invitrogen 10296028) and treated with DNase I (Invitrogen Turbo DNase AM2238). mRNAs were isolated using NEBNext Poly(A) mRNA Magnetic Isolation Module (New England Biolabs E7490) and fragmented using Fragmentation Buffer (Ambion AM8740). cDNAs were synthesized using SuperScript II (Invitrogen 18064014), and non-directional high-throughput sequencing libraries were prepared using NEBNext Ultra DNA Library Prep Kit (New England Biolabs, E7370). Libraries were processed on the Illumina HiSeq platform for single-end 50-nt sequencing. See **Quantification and Statistical Analysis** for downstream analyses. RNA-seq experiments are listed in **Table S3**.

### GRO-seq

Nuclei were isolated by incubating cells in hypotonic NP40 lysis buffer (10 mM NaCl, 3 mM MgCl_2_, 0.5% MP-40, 10 mM Tris, pH 7.5) supplemented with RNase Inhibitor on ice and resuspended in Nuclear Storage buffer (50 mM Tris pH 8.0, 0.1 mM EDTA, 5 mM MgCl_2_, 40% glycerol, RNase inhibitor). The nuclear suspension was mixed with an equal volume of 2x NRO buffer (10 mM Tris pH 8.0, 5 mM MgCl_2_, 1 mM DTT, 300 mM KCl, 0.5 mM ATP, 0.5 mM GTP, 0.5 mM BrUTP, 2 µM CTP). The sample was incubated for 4 min at 30°C without sarkosyl and for 4 min at 30°C with 0.5% sarkosyl (total 8 min). RNAs were purified from the reaction by Trizol LS (Invitrogen 10296028) followed by isopropanol precipitation. RNAs were treated with TurboDNase (Ambion AM18907) and fragmented by Fragmentation Buffer (Ambion AM8740). BrU-incorporated RNA fragments were immunoprecipitated with mouse monoclonal anti-BrdU antibody 3D4 (BD Biosciences 555627 Lot 7033666) and used to construct DNA sequencing libraries using NEBNext Ultra II Directional RNA Library Prep kit (New England Biolabs E7760). DNA libraries were processed on a Illumina NextSeq machine for paired-end 42-nt sequencing. See **Quantification and Statistical Analysis** for downstream analyses. GRO-seq experiments are listed in **Table S3**.

## QUANTIFICATION AND STATISTICAL ANALYSIS

Number of data points, number of replicates, and the type of statistical tests employed are indicated in figures and/or figure legends.

### ELISA quantification

The reaction absorbance (450 nm) minus the reference absorbance (550 nm) was plotted using software R (v3.3.2). Loess fit was computed using *geom_smooth* function in *ggplot2* package (version 2.2.1).

### Western blot quantification

Intensities of Western blot bands and coomassie staining were obtained using the *Analyze Gels* function in Fiji (v1.0). Western blot Intensities were normalized to coomassie staining intensities. The mean and standard deviation of three biological replicates were computed using software R (v3.3.2).

### Flow cytometry quantification

Flow cytometry data were processed using FlowJo (v10.5.3). Forward and side scatter gatings were used to identify single cells. To obtain the fraction of cells at G1/G0, S, and G2/M, DAPI signals were processed with the Watson Pragmatic algorithm (Watson et al., 1987) in FlowJo with manually constrained G1/G0 and G2/M signal ranges with the coefficient of variation set to 10%.

### Immunofluorescence quantification

Quantification of immunofluorescence signals for *LMNA* transgene products was performed on the image at a z-axis position with the highest nuclear periphery-to-interior Lamin A/C signal ratio in a given cell. In this image, a 5-µm line segment that crossed the nuclear periphery at position 0 µm (–2.5 to 2.5 µm with negative coordinates indicating positions outside the nucleus) was drawn. An array of pixel intensities along the segment (averaged across the 10-pixel width) was obtained using Fiji (v1.0). From this array, an average pixel intensity in every 0.1 µm bin along the line was computed (50 bins). To visualize signals along the segment, the binned intensities from the same cell type were quantile normalized using *normalize.quantiles* function in the *preprocessCore* package (v1.36.0) in software R (Bolstad et al., 2003) to account for overall intensity differences between cell types. To compute the nuclear interior-to-periphery ratios, the mean unnormalized signal intensities between +4 to +5 µm of the segment was assigned as the nuclear-interior signal, and the maximum unnormalized signal intensities between –0.5 to +0.5 µm was assigned as the nuclear-interior signal.

### Reference genome

The February 2009 human reference sequence hg19/GRCh37 was used throughout in this paper.

### Blacklisted regions

Before performing genomic data analyses, we excluded all genes and genomic features located in blacklisted regions which are genomic regions that may cause misinterpretation due to high sequence redundancy, uncertain chromosomal locations, high signal background, haplotypes, or potential copy number variations (CNVs) induced in the process of CRISPR-mediated generation of *LMNA^−/−^* cells (Aguirre et al., 2016). The collection of such genomic regions were constructed from the following datasets (see **Data and Code Availability**): assembly gaps in the hg19 reference genome, ENCODE-defined hg19 blacklisted regions, mitochondria sequence (chrM), haplotype chromosomes (chr*_*_hap*), unplaced contigs (chrUn_*), unlocalized contigs (chr*_*_random), and potential CNVs described below. To identify potential large CNVs between wild-type BJ-5ta and BJ-5ta-derived *LMNA^−/−^* cells, input sequencing data for wild-type BJ-5ta (ID KI481) and *LMNA^−/−^* cells (ID KI489) were processed with CNV-seq (Xie and Tammi, 2009) with the following parameters: [–genome human –global-normalization –log2-threshold 0.5 –minimum-windows-required 3]. After removing windows with low sequence coverage, candidate CNV windows that were overlapping or spaced within 500 kb were merged, and isolated windows smaller than 500 kb in size were removed. This yielded 5 candidate large CNVs (24 Mb in chr1; 15 Mb in chr 4; 2.7 Mb in chr19; 683 kb in chr2; and 528 kb in chrX). The blacklisted regions are listed in **Table S4**.

### Gene annotation

The Gencode V19 “Basic” gene annotation was downloaded from the UCSC genome browser. Of the total 99,901 transcripts in the list, we retained transcripts that met all of the following requirements: 1) “gene type” equals “transcript type”; 2) “transcript type” is either *protein_coding* or *antisense* or *lincRNA*; and 3) “transcript ID” appears only once in the list. This processing yielded 75,968 transcripts. To select one transcribed unit per gene locus, the 75,968 transcripts were grouped by “gene symbol,” and within the group, transcripts were sorted by the “exon count” (largest first), then by “length” of transcribed region (largest first), then by the alphanumeric order of the “transcript ID” (smallest first), and the transcript that appeared first in the group was chosen to represent the transcribed unit of that gene. In this processing, in general, a transcript with a largest number of annotated exons among other associated transcripts represented the gene. After removing genes located within the blacklisted regions (see *Blacklisted regions*), we obtained 31,561 genes, which included 19,469 “protein_coding” genes.

### ChIP-seq data processing

ChIP-seq experiments and sequencing depth are listed in **Table S3**. ChIP-seq reads were mapped to the hg19 human reference genome using Bowtie2 (Langmead and Salzberg, 2012) with the default “--sensitive” parameter. Reads with MAPQ score greater than 20 were used in downstream analyses. Reads from biological replicates of ChIP and the corresponding input were processed by MACS2 (v2.1.0) (Zhang et al., 2008). MACS2 removed duplicated reads and generated then “fold enrichment score”, which was input-normalized per-base coverage of 200 nt-extended reads. We used the fold-enrichment scores throughout the paper for quantitative analysis of ChIP-seq enrichment. In addition, for ChIP-seq with point-source enrichment profiles, MACS2 was used to identify statistically overrepresented peak regions and peak summits using the following parameter set: [call peak -g hg --nomodel --extsize 200 -- call-summits]. Peaks overlapping blacklisted regions (see *Blacklisted regions* section) were removed. We defined a summit as a unit of a protein binding site. MACS2 identified 22,966 pS22-Lamin A/C-binding sites in BJ-5ta (**Table S5**); 79,799 H3K27ac-enriched sites in BJ-5ta (**Table S8**); 18,100 H3K4me3-enriched sites in BJ-5ta (**Table S9**); 87,988 c-Jun-enriched sites in BJ-5ta (**Table S10**). The codes used are publicly available (see **Data and Code Availability**).

### ATAC-seq data processing

ATAC-seq experiments and sequencing depth are listed in **Table S3**. For alignment, the first 38 nt of the 41-nt reads in the 5′ to 3′ direction were used. The rationale of this trimming is that the minimum size of DNA fragments that can be flanked by Tn5 transposition events has been estimated to be 38 bp (Reznikoff, 2008; Picelli et al., 2014), and therefore, a 41-nt read could contain a part of read-through adaptors. We aligned 38-nt reads to the hg19 reference genome using bowtie2 with following parameters: [-X 2000 --no-mixed --no-discordant --trim3 3]. The center of the active Tn5 dimer is estimated to be located +4-5 bases offset from the 5′-end of the transposition sites (Reznikoff, 2008; Picelli et al., 2014). To place the Tn5 loading center at the center of aligned reads, the 5′-end of the plus-strand read was shifted 4 bp in the 5′-to-3′ direction and that of the minus-strand reads was shifted 5 bp in the 5′-to-3′ direction, and the shifted end (1 nt) was extended +/–100 bp. To generate background datasets that capture local bias of read coverage, the shifted read ends were extended +/–5,000 bp and used to construct local lambda background file. We processed the Tn5 density file and the local lambda file (background) with MACS2 function *bdgcomp* to generate the fold-enrichment scores for Tn5 density from as a background. We also used MACS2 function *bdgpeakcall* to identify regions with statistically significant Tn5 enrichment (ATAC peaks). At P-value cutoff of 1×10^-10^, we obtained 73,933 ATAC peaks (**Table S7**). The codes used are publicly available (see **Data and Code Availability**).

### RNA-seq data processing

RNA-seq experiments and sequencing depth are listed in **Table S3**. Public RNA-seq raw data files (fastq) for normal and progeria-patient fibroblasts (Fleischer et al., 2018) (see **Data and Code Availability**) were retrieved using *fastq-dump* (version 2.9.3). All RNA-seq reads were aligned to the hg19 human reference genome using Tophat2 with the default parameter set (Kim et al., 2013). Reads with MAPQ score greater than 50 were used in downstream analyses. For each of the total 31,561 genes (see *Gene annotation*) for each replicate, we computed (1) unnormalized RNA-seq read coverage, which was the sum of per-base read coverage in exons; and (2) RPKM (Reads Per Kilobase of transcript per Million mapped reads), which was the unnormalized RNA-seq read coverage divided by the read length (50 nt) and then by the sum of the exon size in kilobase and then by the total number of reads in million. The RPKM scores were Log_2_-transformed (Log_2_(RPKM+0.001)), z-normalized (for each sample), and then used in Principal Component Analysis (PCA) using *prcomp* function (*stats* package version 3.3.2) in R. The PCA found that two data sets, s78 (normal fibroblast GM05381 by Fleischer et al.) and KI429 (normal fibroblast GM08398 by this study), did not cluster with other normal-fibroblast datasets (**Fig. S4D**). A retrospective assessment of the RNA quality of KI429 found a sign of RNA degradation. The datasets s78 and KI429 were excluded from the subsequent analyses. The codes used are publicly available (see **Data and Code Availability**).

### GRO-seq data processing

GRO-seq experiments and sequencing depth are listed in **Table S3**. GRO-seq read pairs with maximum fragment length of 2000 bp were aligned to the hg19 human reference genome using Bowtie2 with the parameter set of *-X 2000 --no-mixed --no-discordant*. Reads with MAPQ score greater than 20 were used in downstream analyses. For each of the total 31,561 genes (see *Gene annotation*), we computed “normalized GRO-seq base coverage,” which was the sum of GRO-seq fragment per-base coverage normalized to gene lengths and sequencing depths. The codes used are publicly available (see **Data and Code Availability**).

### LADs

LADs in BJ-5ta were defined using pan-N-terminal-Lamin A/C ChIP-seq data in BJ-5ta. The hg19 genome was segmented into 5-kb non-overlapping windows, and for each window, the sum of perbase read coverage (from replicate-combined reads) was computed for pan-N-terminal-Lamin A/C ChIP-seq and the corresponding input. The coverage was normalized by sequencing depth. The depth-normalized coverage was used to compute per-window log_2_ ratios of ChIP over input. We then created 100 kb windows with a 5-kb step genome-wide. For each 100-kb window, if every one of the 20 constituting 5-kb windows had a positive log_2_ ratio and the mean log_2_ ratios of the constituting 5-kb windows was greater than 0.5, this 100-kb window was further processed. The qualified 100-kb windows were merged if overlapping or touching. After filtering regions overlapping blacklisted regions (see *Blacklisted regions*), we obtained 2,178 regions which we defined as Lamin A/C LADs in BJ-5ta (**Table S6**). LADs defined by Lamin B1 Dam ID in lung fibroblasts are reported previously (Guelen et al., 2008) (see **Data and Code Availability**).

### Analysis on deciles

Genes, ATAC-seq sites, or c-Jun-binding sites were stratified into deciles by scores specified in figure legends using *ntile* function in dplyr (version 0.7.4) in R.

### DNA motif analysis

To find DNA motifs *de novo*, 150-bp sequences centered around the summit of the top 500 high-confident (defined by p-values) pS22-Lamin A/C-binding sites were analyzed by MEME (v4.10.0) (Machanick and Bailey, 2011) with the following parameters: minimum motif size, 6 bp; maximum motif size, 12 bp; and the expected motif occurrence of zero or one per sequence (*-mod zoops*) and with the 1st-order Markov model (i.e. the dinucleotide frequency) derived from the 73,933 ATAC-seq sites as the background. The top five overrepresented motifs were then processed by TOMTOM (v4.11.3) (Gupta et al., 2007; Tanaka et al., 2011) to identify known motifs that corresponded to the *de novo* identified motifs from human HOCOMOCOv10 motif database (Kulakovskiy et al., 2018). At the q-value cutoff of less than 0.001, the motifs #1 (AP1), #2 (FOX) and #4 (RUNX) found the corresponding known motifs in the database. The location and frequency of these motifs within the all pS22-Lamin A/C-binding sites (total 22,966) were determined by FIMO (v4.10.0) with p-value threshold less than 0.0001.

### Upregulated and downregulated genes in progeria

To identify differentially expressed genes between progeria-patient fibroblasts and normal-individual fibroblasts, we applied DESeq2 (version 1.14.1) (Love et al., 2014) on the unnormalized RNA-seq read coverage for the total 31,561 genes (see *RNA-seq data processing*). To account for the variables due to the study origins, we included the study origin annotation as an additive term in the DESeq2 model. We applied the cutoff of DESeq2-computed adjusted p-value smaller than 0.05 and absolute DESeq2-adjusted log_2_-fold change greater than 0.5. Among 11,613 protein coding genes that had the minimum RPKM score greater than 0.01 across all analyzed samples (“expressed genes”), 615 genes were defined as upregulated and 502 genes were defined as downregulated in progeria-patient fibroblasts compared with normal-individual fibroblasts (**Table S11**). The codes used are publicly available (see **Data and Code Availability**).

### Gained and lost pS22-Lamin A/C-binding sites in progeria

We first generated a union set of pS22-Lamin A/C-binding sites in normal-individual fibroblast GM07492 and progeria-patient fibroblast AG11498 by collecting pS22-Lamin A/C-enriched sites from the two cell lines and then merging neighboring summits if the distance between the summits was equal to or smaller than 200 bp. When summits were merged, the center of the region generated by the merged summits was assigned as the new summit. This resulted in 15323 union pS22-Lamin A/C-binding site summits (**Table S12**). The summits were then extended +/–500 bp, and the sum of per-base read coverage of pS22-Lamin A/C ChIP-seq (reads extended to 200 bp) within the 1000-bp regions was computed for each replicate of the normal and progeria fibroblasts (two replicates each). This coverage matrix was processed using DESeq2 (version 1.14.1) (Love et al., 2014) to identify pS22-Lamin A/C-binding sites with statistically-significant difference between the normal and progeria-patient fibroblast cell lines. At DESeq2-computed adjusted p-value < 0.05 and absolute log_2_-fold change > 0.5, we identified 2,796 sites whose pS22-Lamin A/C signals were higher in progeria fibroblasts than in normal fibroblasts (gained pS22-Lamin A/C-binding sites) and 2,425 sites whose pS22-Lamin A/C signals were higher in normal fibroblasts than in progeria fibroblasts (lost pS22-Lamin A/C-binding sites) (**Table S12**).

### Box plot and Violin plot

Throughout the paper, box plots with or without overlaid “violins” were used to visualize distribution of numeric data. The box indicates interquartile (IQR) range with the bar inside the box indicating the median. The upper and lower whiskers indicate maximum and minimum data points within 1.5x IQR from the box, respectively. The circles outside the whisker range indicate data points outside of this range. The overlaid violin indicates the kernel density of the data. Box and violin plots were generated using *ggplot2* package (version 2.2.1) in R (version 3.2.2).

### Gained and lost LADs in progeria

We first generated 5-kb windows genome-wide, and for each window and for each cell type (normal-individual GM07492 and progeria-fibroblast AG11498), we computed the mean of the replicate-combined pan-N-terminal-Lamin A/C ChIP-seq log2-fold-enrichment scores within the window. The data was then quantile-normalized using *normalize.quantiles* function in the *preprocessCore* package (v1.36.0) in software R (Bolstad et al., 2003). From the normalized data, we identified: (1) “seeds” of lost LADs, which were 5-kb windows whose normal-fibroblast score was a positive number and whose progeria-fibroblast score was a negative number; (2) “seeds” for gained LADs, which were 5-kb windows whose normal-fibroblast score was a negative number and whose progeria-fibroblast score was a positive number; and (3) “seeds” for steady LADs, which were 5-kb windows whose normal-fibroblast and progeria-fibroblast scores were both positive. For gained and lost LADs, the neighboring aforementioned seeds were merged (individually for gained and lost LAD seeds) if they were located within 10 kb. For steady LADs, seeds were merged if the neighboring seeds were located within 5 kb. Finally, the merged windows greater than 100 kb in size were isolated, and then those intersecting blacklisted regions (see *Blacklisted regions*) were removed. These processing resulted in 282 gained LADs, 353 lost LADs, and 2,100 steady LADs (**Table S13**).

### Aggregate plot and heatmap

To generate aggregate plots and heatmaps for features, two data files were first generated: (a) a window file, which consists of, for each genomic feature, a set of fixed-size genomic windows that cover genomic intervals around the feature; and (b) a genome-wide signal file (in bedgraph format). For each genomic window in the window file, all signals within that window were obtained from the signal file, and either mean or max of the signals or sum of the perbase signal (“area”) were computed.

For the heatmaps and aggregate plots around pS22-Lamin A/C-binding sites or ATAC sites, a set of 250-bp windows with a 50-bp offset that covered a 10-kb region centered around the summit of these sites was generated for each site. For each window, the mean of fold-enrichment score was computed from replicate-combined input-normalized fold enrichment bedgraph files.

For the heatmaps of LADs, a set of 5-kb windows (without an offset) that covered the LAD body or the LAD body plus 250 kb downstream region was generated for each LAD. For each window, the mean of fold-enrichment score was computed from replicate-combined input-normalized fold enrichment bedgraph files.

For the heatmap of the differentially expressed genes, RPKM scores for the 11,613 expressed genes for the 26 RNA-seq data sets (12 normal fibroblasts and 14 progeria-patient fibroblasts) were log_10_-transformed and quantile-normalized using *normalize.quantiles* function in the *preprocessCore* package (v1.36.0) in R (Bolstad et al., 2003). These normalized scores were then processed using *ComBat* function in the *sva* package (v3.22.0) in R with the sample origin annotation as the batch to remove (Johnson et al., 2007). The batch-normalized RPKM scores for the total 1,117 dysregulated genes in progeria fibroblasts were clustered and visualized using *pheatmap* function in the *pheatmap* package (v1.0.12) in R with the “correlation” clustering distance measurement method between sample clusters.

The computed scores were plotted using *ggplot2* package (version 2.2.1) in R (version 3.2.2). The codes used are publicly available (see **Data and Code Availability**).

### Gene ontology analysis

Gene ontology (GO) analyses were performed using Metascape (Tripathi et al., 2015). The input data type was Gencode 19 *Gene Symbol* (see *Gene annotation*). For background, the 11,613 protein-coding genes with reliable sequencing coverage were used (see *Differentially expressed genes*). Enrichment for “DisGeNet” terms (Piñero et al., 2017) was analyzed under the default parameter settings (minimum gene count 3, P <0.01, enrichment over background >1.5).

### One-way ANOVA

One-way ANOVA was used to assess statistical difference of numeric scores among cells expressing wild-type or phospho-mutant Lamin A/C (6 cell types). The ANOVA tests were performed using *aov* function with default parameters in R (*stats* package v3.3.2) under the null hypothesis that the population mean of the numeric scores was the same across all cell types, with the alternative hypothesis that at least one cell type had a different population mean. We then applied post-hoc Tukey’s test to perform pairwise comparison. The Tukey’s test was performed using *TukeyHSD* function on the ANOVA result above with default parameters in R (*stats* package v3.3.2) under the null hypothesis that the population mean of the numeric scores between the two cell types being compared was the same, with the alternative hypothesis that they were different.

### Mann-Whitney U test

Throughout the paper, Mann-Whitney *U* test was used to assess statistical difference of numeric scores between two groups. The tests were performed using *wilcox.test* function with default parameters in R (*stats* package v3.3.2) under the null hypothesis that numeric scores of the two groups were selected from one population, with the alternative hypothesis that they came from different populations.

### Fisher’s exact test

Throughout the paper, Fisher’s exact tests were used to assess the association between two features that were unambiguously and independently assigned to each data point in one data set. The tests were performed on a 2-by-2 contingency table using *fisher.test* function with default parameters in R (*stats* package v3.3.2) under the null hypothesis that the odds ratio is equal to 1, with alternative hypothesis that the odds ratio is not equal to 1.

### One-sample Student’s t test

One-sample *t*-tests were performed using t.test function with default parameters in R (stats package v3.3.2) under the null hypothesis that the mean of the numeric vector equal to zero, with the alternative hypothesis that the mean is not equal to zero.

### Permutation test for chromatin state analysis

For the 22,966 pS22-Lamin A/C-binding site summits (1 bp), a random set of 22,966 genomic locations was selected from the blacklisted-region-filtered genome using *shuffle* function in Bedtools (v2.25.0) (Quinlan and Hall, 2010) such that the chromosome distribution of the original 22,966 pS22-Lamin A/C-binding sites was maintained. This process was iterated 2,000 times. For each iteration, the total number of bases overlapped with a given chromatin state was computed. For each chromatin state, the number of iterations in which the base coverage of permutated pS22-Lamin A/C-binding sites exceeded the actual base coverage of the 22,966 pS22-Lamin A/C-binding sites was counted. If this number is 0 (one-sided test for over-representation), we assigned empirical P-value of < 0.001. Essentially, the same computation was performed for 2,178 LADs, except that the random sets of LADs maintained the distribution of the original feature sizes.

### Correlation analysis

Pearson correlation coefficient was used to assess correlation between two fold-enrichment scores at 22,966 pS22-Lamin A/C-binding sites (sum of fold-enrichment scores within +/–500 bp of the site center). The computation of Pearson’s *r* was performed using *cor* function in R with *method=”pearson”* (*stats* package v3.3.2).

Spearman’s rank correlation coefficient was used to assess the monotonic relationship between gene deciles by GRO-seq coverage and the fraction of genes linked to pS22-Lamin A/C-binding sites within each of the deciles. The computation of Spearman’s *rho* was performed using *cor* function in R with *method=”spearman”* (*stats* package v3.3.2).

### GO-term enrichment

P-values indicating GO-term enrichment were computed using Metascape (Tripathi et al., 2015), which uses a cumulative hypergeometric statistical test.

## DATA AND CODE AVAILABILITY

### Data and Code Availability Statement

The genomic datasets generated during this study are available at Gene Expression Omnibus (http://www.ncbi.nlm.nih.gov/geo/) under the accession number GSE113354.

Custom computational scripts used in this study are available at: https://github.com/kohta-ikegami/pS22-LMNA/

### Public datasets

Assembly gaps http://hgdownload.soe.ucsc.edu/goldenPath/hg19/database/gap.txt.gz

Blacklisted Regions http://hgdownload.cse.ucsc.edu/goldenPath/hg19/encodeDCC/wgEncodeMapability/wgEncodeDacMapabilityConsensusExcludable.bed.gz

Chromatin annotation http://egg2.wustl.edu/roadmap/data/byFileType/chromhmmSegmentations/ChmmModels/coreMarks/jointModel/final/E126_15_coreMarks_stateno.bed.gz

Genecode Release 19 Gene Model http://hgdownload.cse.ucsc.edu/goldenPath/hg19/database/wgEncodeGencodeBasicV19.txt.gz

Lamin B1 LADs http://hgdownload.cse.ucsc.edu/goldenPath/hg19/database/laminB1Lads.txt.gz

Progeria fibroblast RNA-seq datasets from Fleischer et al. https://www.ncbi.nlm.nih.gov/geo/query/acc.cgi?acc=GSE113957

## SUPPLEMENTARY TABLES

All Supplementary Tables are presented in the tabs in the Excel spreadsheet.

### Excel file Set1

Table S1: Oligonucleotide sequences (Text)

Table S2: Normal and progeria fibroblast cell lines (Text)

Table S3: High-throughput sequencing data sets (Text)

Table S4: Blacklisted regions (UCSC Bed)

### Excel file Set2

Table S5: pS22-Lamin A/C-binding sites in BJ-5ta (UCSC Bed)

Table S6: LADs in BJ-5ta (UCSC Bed)

Table S7: ATAC-seq peaks in BJ-5ta (UCSC Bed) Excel file Set3:

Table S8: H3K27ac ChIP-seq peaks in BJ-5ta (UCSC Bed)

Table S9: H3K4me3 ChIP-seq peaks in BJ-5ta (UCSC Bed)

Table S10: c-Jun ChIP-seq peaks in BJ-5ta (UCSC Bed) Excel file Set4:

Table S11: Differentially-expressed genes in progeria (UCSC Bed)

Table S12: Gained and lost pS22-Lamin A/C-binding sites in progeria (UCSC Bed)

Table S13: Gained and lost LADs in progeria (UCSC Bed)

Table S14: Progeria-up genes linked to gained pS22-Lamin A/C-binding sites with DisGeNet annotation (Text)

---

**Figure S1.**
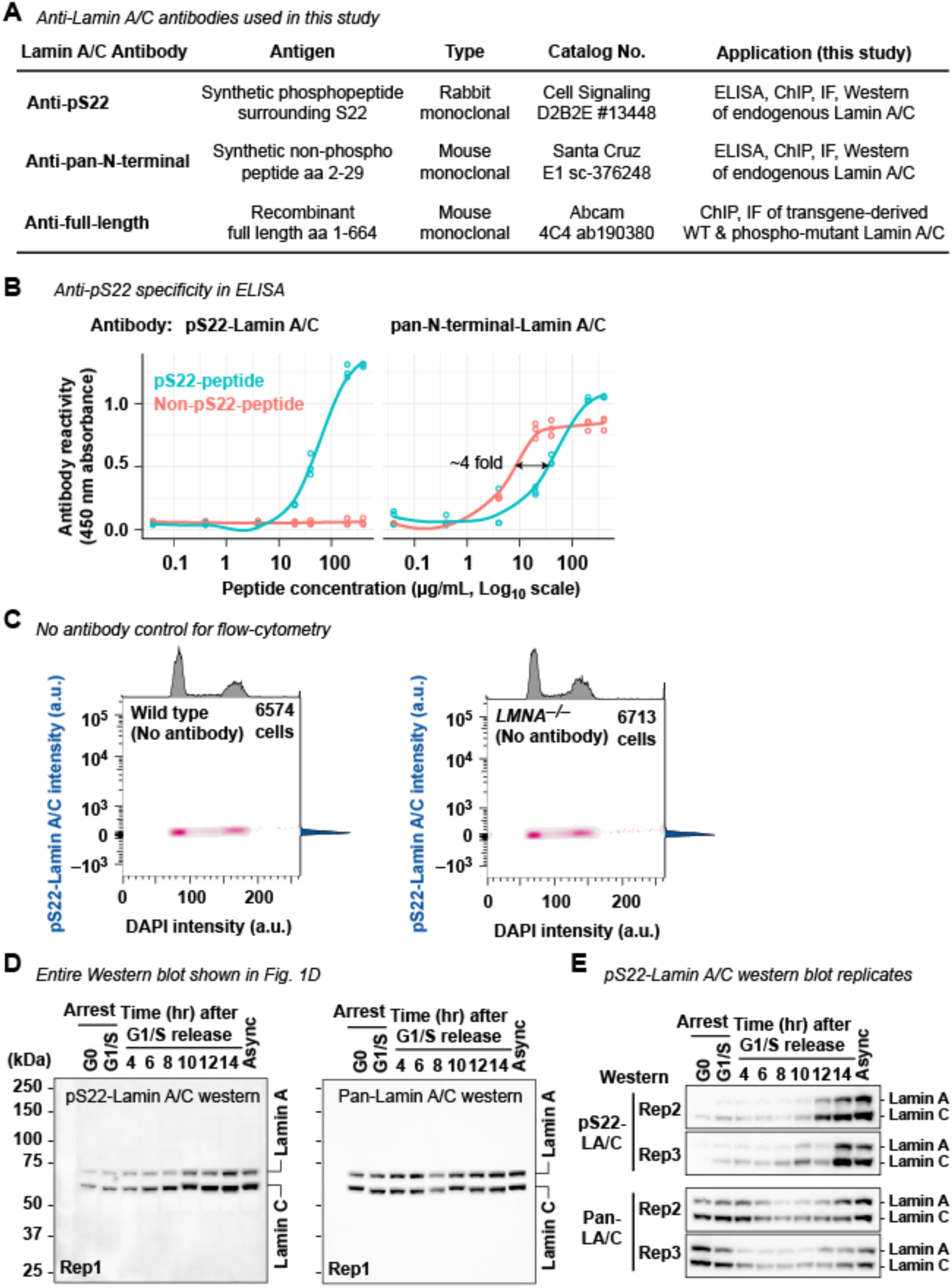
Characterization of S22-phosphorylated Lamin A/C (related to Figure 1) **(A)** Anti-phospho-S22-Lamin A/C antibody and anti-pan-N-terminal-Lamin A/C antibody used in this study. **(B)** ELISA on immobilized synthetic phospho-S22 or non-phospho-S22-Lamin A/C peptides (aa2-29) incubated with anti-pS22-Lamin A/C or anti-pan-N-terminal-Lamin A/C antibodies. Circle, technical replicate. Line, Loess fit. **(C)** Control flow cytometry analysis without anti-pS22-Lamin A/C antibody for **Fig. 1C**. **(D)** Uncropped images of Western blots for synchronized wild-type BJ-5ta, part of which are shown in **Fig. 1D**. **(E)** (Top) Same as **D**, but two additional biological replicates are shown.

**Figure S2.**
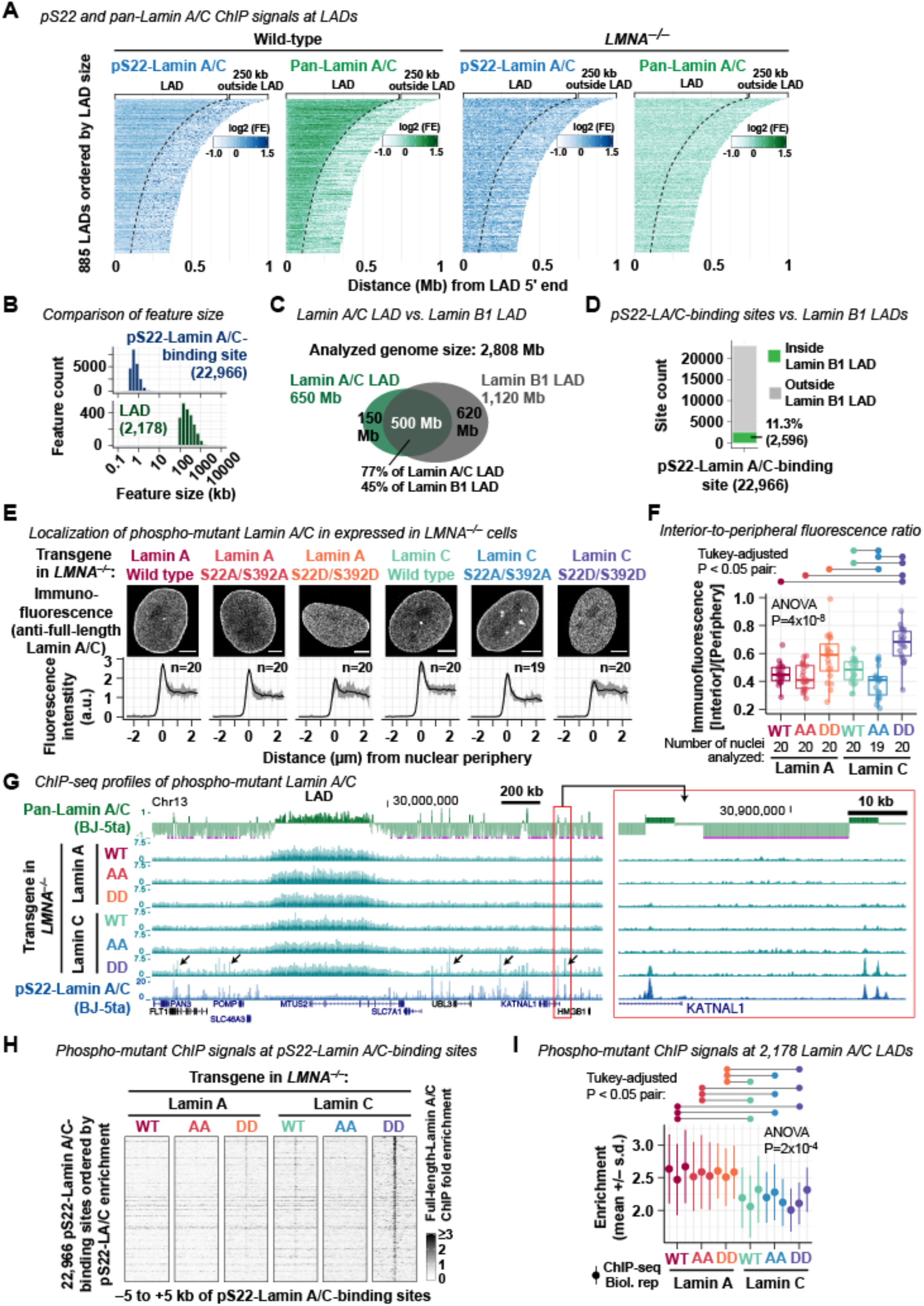
Characterization of pS22-Lamin A/C-binding sites (related to Figure 2) **(A)** ChIP-seq fold-enrichment (FE) scores at Lamin A/C LADs defined by pan-N-terminal-Lamin A/C ChIP-seq. A subset of LADs that do not have adjacent LADs within 250 kb downstream of LADs are shown. **(B)** Histogram of the size of pS22-Lamin A/C-binding sites and Lamin A/C LADs. **(C)** Comparison of genomic domains covered by Lamin A/C LADs (in BJ-5ta, this study) or Lamin B1 LADs (lung fibroblast, Guelen et al., 2008). **(D)** Location of pS22-Lamin A/C-binding sites with respect to the 1,302 Lamin B1 LADs. **(E)** (Top) Immunofluorescence of transgene-driven Lamin A or Lamin C with phospho-deficient S22A/S392 mutations, or phospho-mimetic S22D/S392D mutations, or without mutation, expressed in *LMNA^−/−^* cells. Anti-full-length-Lamin A/C antibody was used. (Bottom) Immunofluorescence signal distribution along a 5-µm segment that crosses the nuclear periphery at 0 µm (–2.5 to 2.5 µm with negative coordinates indicating positions outside the nucleus). Line, mean. Shade, minimum to maximum. n, number of nuclei analyzed. **(F)** Interior-to-periphery Lamin A/C immunofluorescence signal ratio in transgene-driven wild-type (WT), phospho-deficient (AA), and phospho-deficient (DD) Lamin A/C expressed in *LMNA^−/−^* cells. Interior signals are the mean signal between +4 to +5 µm of an aforementioned 5-µm segment. Peripheral signals are the maximum signal between -0.5 to +0.5 µm of the segment. One-way ANOVA compares the ratio of all isoforms, with post-hoc Tukey analysis for pairwise comparison (pairs with P<0.05 are indicated). **(G)** Representative ChIP-seq profiles for transgene-driven wild-type (WT), phospho-deficient (AA), and phospho-deficient (DD) Lamin A/C expressed in *LMNA^−/−^* cells. Anti-full-length-Lamin A/C antibody was used in ChIP. Signals are FE scores. Pan-N-terminal Lamin A/C and pS22-Lamin A/C ChIP-seq profiles in BJ-5ta are shown for comparison. Arrows indicate positions at which phospho-mimetic Lamin C was enriched at pS22-Lamin A/C binding sites. **(H)** ChIP-seq FE scores at the 22,966 pS22-Lamin A/C-binding sites. **(I)** Mean ChIP-seq FE scores across the 2,178 Lamin A/C LADs for each biological replicate. Mean of FE scores in each LAD is averaged across all LADs. One-way ANOVA compares means of all isoforms, with post-hoc Tukey analysis for pairwise comparison (pairs with P<0.05 are indicated).

**Figure S3.**
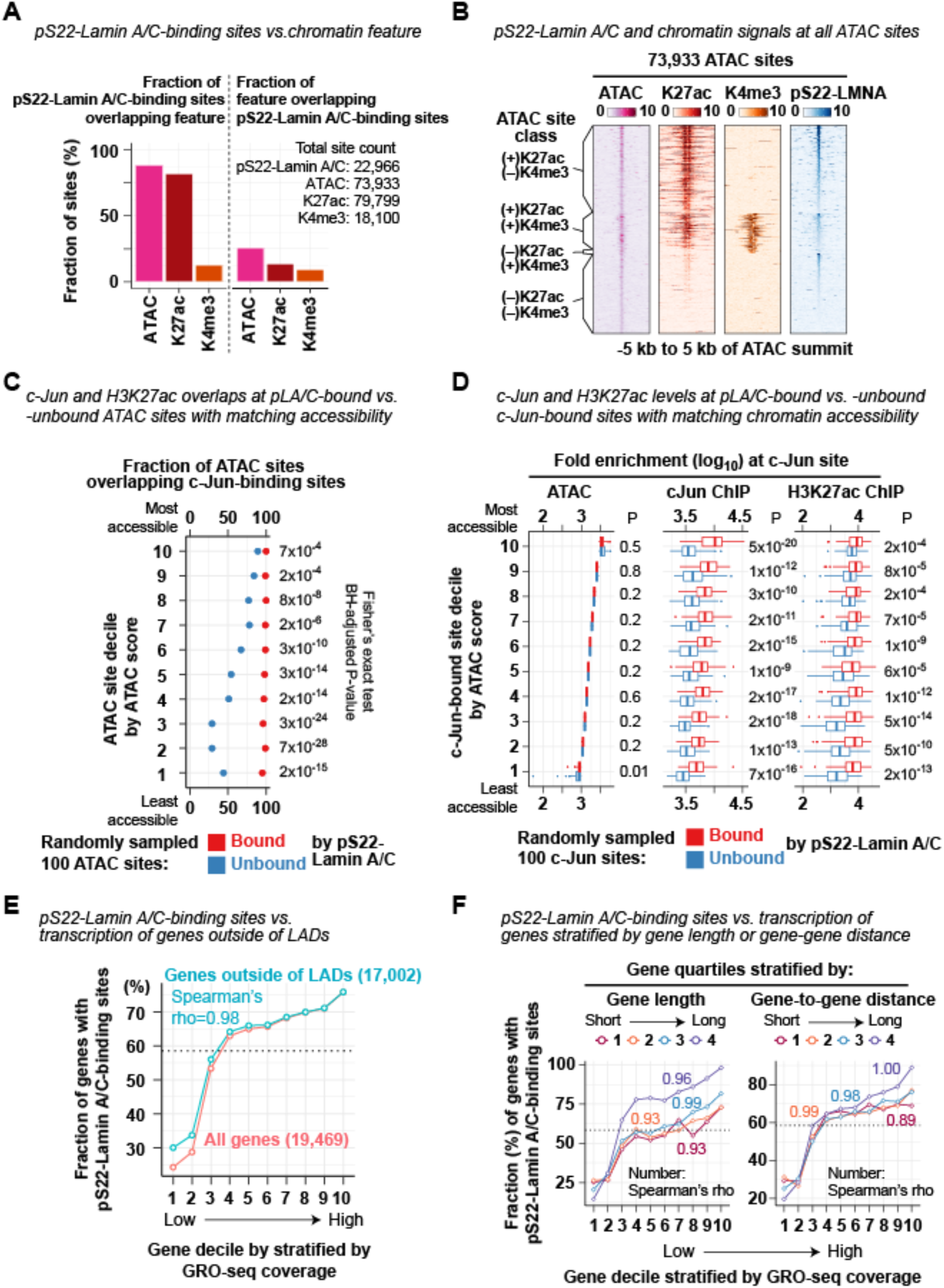
Chromatin characteristics of pS22-Lamin A/C-binding sites (related to Figures 2 and 3) **(A)** Fraction of the 22,966 pS22-Lamin A/C-binding sites that overlap ATAC-seq-defined accessible sites, H3K27ac ChIP-seq peaks, or H3K4me3 ChIP-seq peaks in BJ-5ta. **(B)** ATAC-seq and ChIP-seq fold-enrichment (FE) scores at the 73,933 ATAC-seq-defined accessible sites in BJ-5ta. **(C)** Fraction of pS22-Lamin A/C-bound or -unbound ATAC sites overlapping c-Jun-binding sites. ATAC sites (total 73,933 sites) are stratified into deciles by ATAC FE scores, and from each decile, 100 pS22-Lamin A/C-bound and 100 pS22-Lamin A/C-unbound ATAC sites are randomly selected, and the intersection is assessed. Fisher’s exact test compares association between being bound by pS22-Lamin A/C and being bound by c-Jun, and P-values are adjusted by the Benjamini-Hochberg method. **(D)** c-Jun and H3K27ac ChIP-seq FE scores at c-Jun-binding sites bound or unbound by pS22-Lamin A/C. c-Jun-binding sites (total 87,988 sites) are stratified into deciles by ATAC FE scores, and from each decile, 100 pS22-Lamin A/C-bound and 100 pS22-Lamin A/C-unbound ATAC sites are randomly selected, and their FE score distribution is visualized. Sum of FE scores within +/–250 bp of ATAC site center is used for analysis. Mann-Whitney *U*-test compares FE scores between pS22-Lamin A/C-bound and -unbound sites, and P-values are adjusted for multiple comparisons by the Benjamini-Hochberg method. **(E)** Fraction of genes with pS22-Lamin A/C-binding sites (within gene body or 100 kb upstream) in gene decile stratified by GRO-seq coverage in BJ-5ta. Genes are first stratified by GRO-seq coverage and then grouped by their TSS positions with respect to Lamin A/C LADs (either overlapped or not). Data for all genes (shown in **Fig. 3H**) is shown again here for reference. Dotted line, fraction of all genes with pS22-Lamin A/C-binding sites. **(F)** Same as **(E)**, but genes are stratified (after GRO-seq stratification) by gene length (left) or distance to closest genes (right).

**Figure S4.**
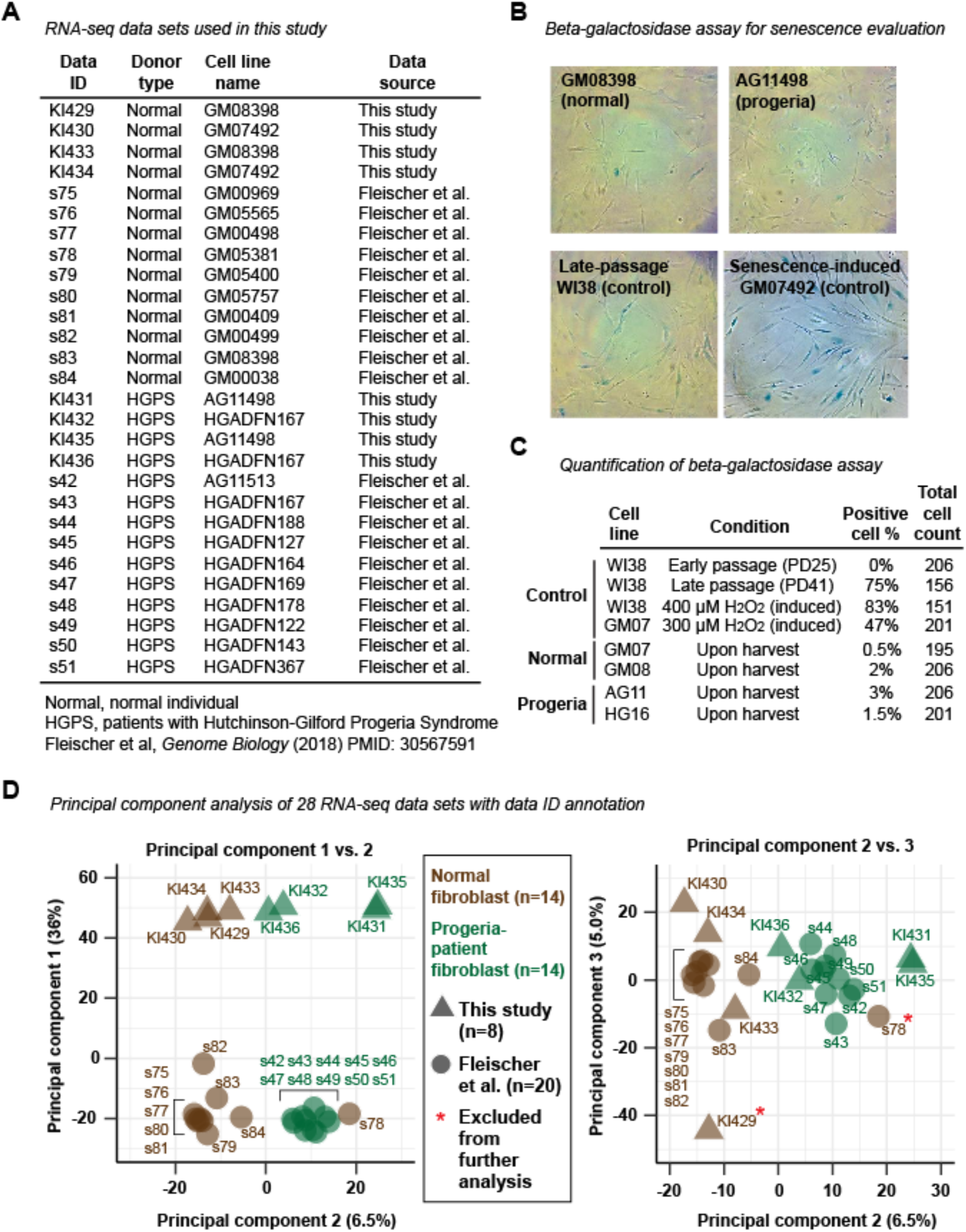
RNA-seq in progeria-patient fibroblasts (related to Figure 4) **(A)** The 28 RNA-seq datasets used in this study. **(B)** Representative images of beta-galactosidase assay evaluating the extent of cell senescence. The assay was performed in the first biological replicate of cells used in RNA-seq and ChIP-seq. **(C)** Quantification of beta-galactosidase assay. Early-passage WI38 fibroblasts serve as a negative control. Late-passage WI38 serves as a positive control. WI38 and GM07492 cells treated with H_2_O_2_ serve as additional positive controls. Cell line notation: GM08, GM08398. GM07, GM07492. AG11, AG11498. HG16, HGADFN167. **(D)** Principal component analyses on RNA-seq RPKM for all 31,561 genes (see *Gene annotation* in **Methods**). The left panel without data ID annotation is shown in **Fig. 4A**.

**Figure S5.**
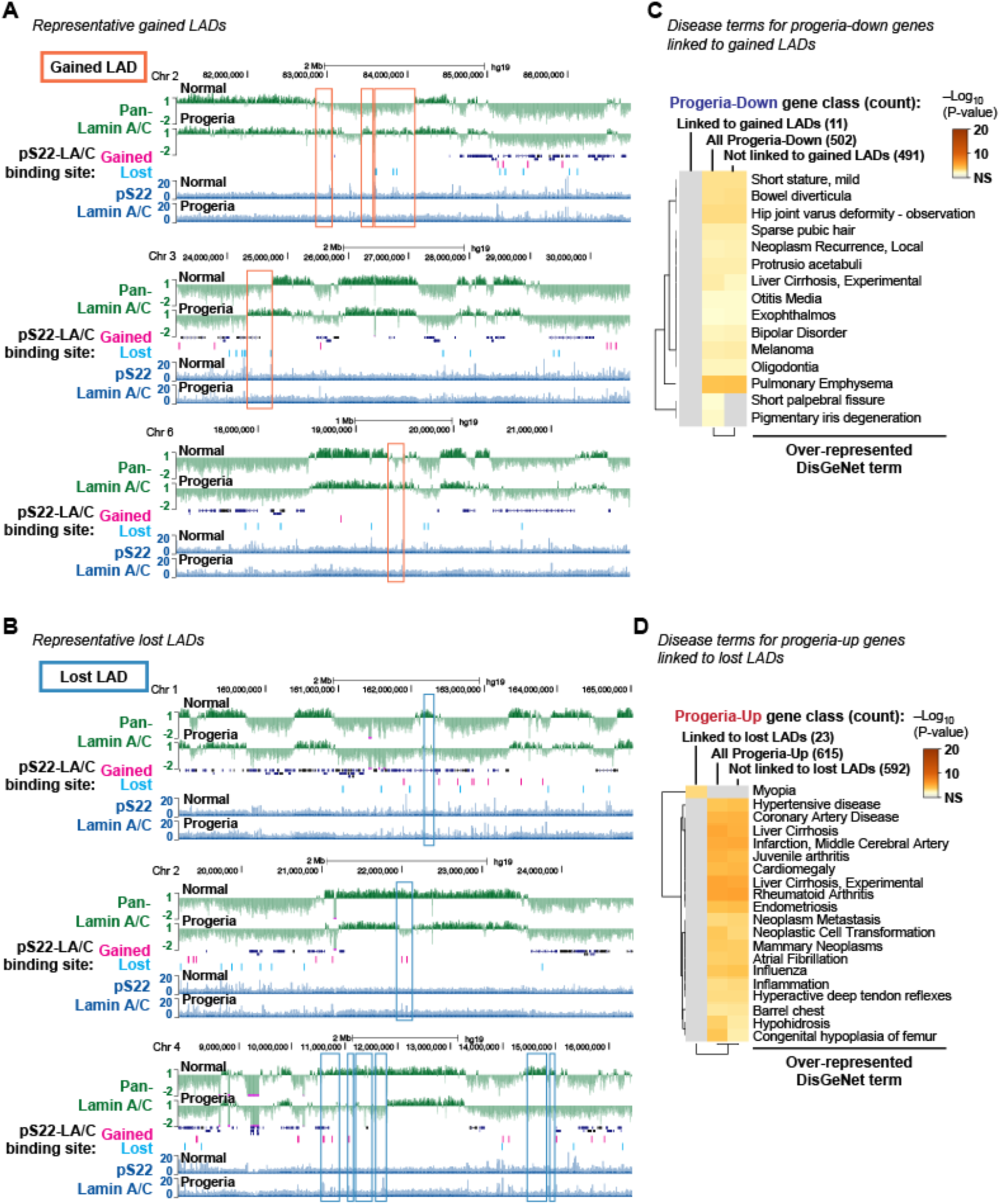
Gained and lost LADs in progeria-patient fibroblasts (related to Figure 5) **(A)** Representative pan-N-terminal-Lamin A/C (pan-Lamin A/C) and pS22-Lamin A/C ChIP-seq profiles in normal (GM07492) and progeria-patient (AG11498) fibroblasts. Signals are fold-enrichment (FE) scores. Rectangle, gained LADs in progeria-patient fibroblasts. Pan-N-terminal-Lamin A/C profiles are in the log_2_ scale to visualize LAD patterns. **(B)** Same as **(A)**, but rectangles indicate lost LADs in progeria-patient fibroblasts. **(C)** DisGeNet-curated disease terms over-represented among progeria-down genes linked to gained LADs, progeria-down genes not linked to gained LADs, and all progeria-down genes. No terms are enriched among progeria-down genes linked to gained LADs. **(D)** Same as **(C)**, but progeria-up genes linked to lost LADs, progeria-up genes not linked to lost LADs, and all progeria-up genes are shown.

**Figure S6.**
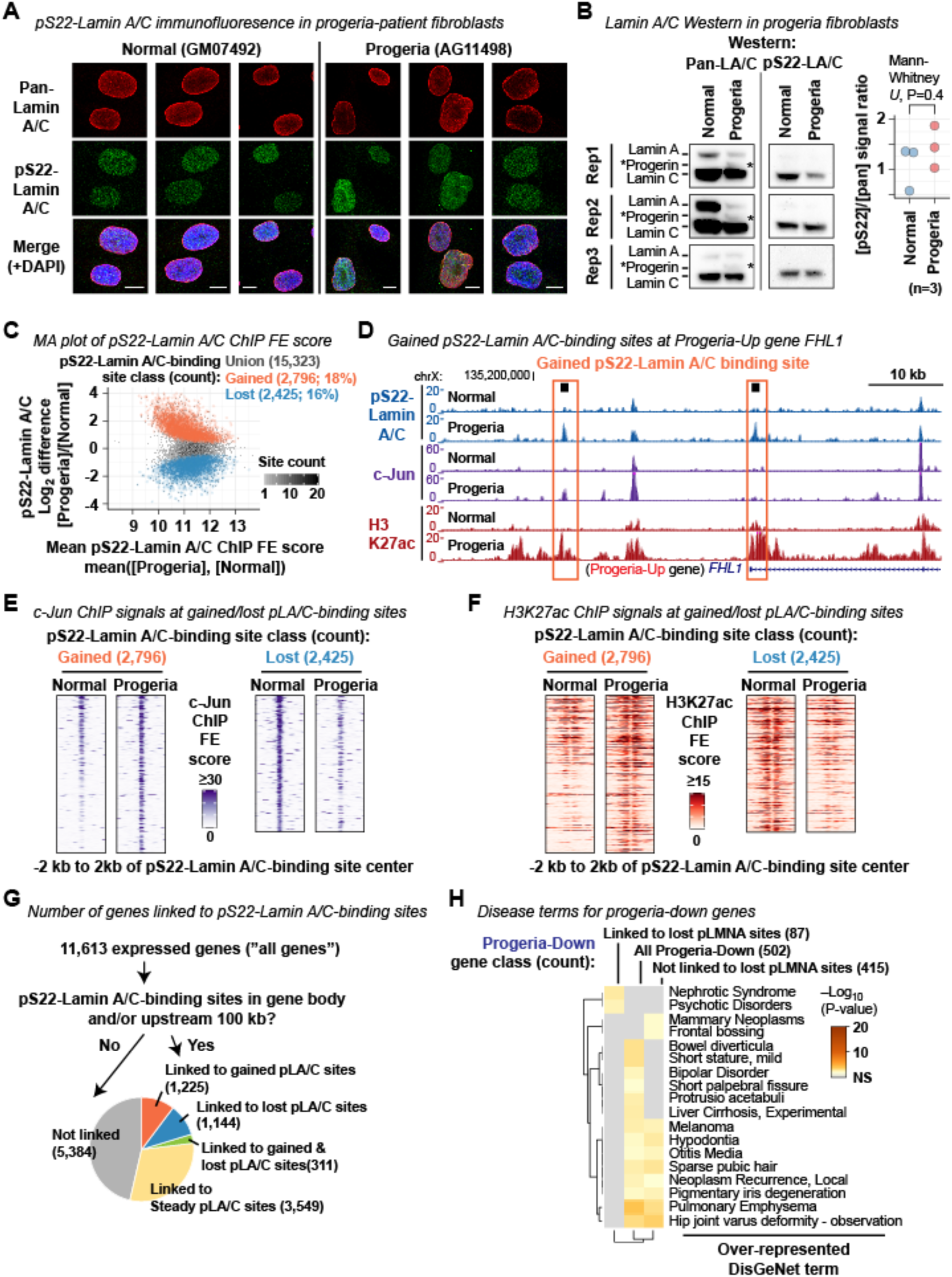
Gained and lost pS22-Lamin A/C-binding sites in progeria-patient fibroblasts (related to Figure 6) **(A)** Immunofluorescence using anti-pan-N-terminal Lamin A/C antibody and anti-pS22-Lamin A/C antibody in normal and progeria fibroblasts. Representative images from 3 biological replicates are shown. **(B)** (Left) Western blot for pan-N-terminal-Lamin A/C and pS22-Lamin A/C on normal (GM07492) and progeria fibroblast (AG11498) extracts. Asterisk, progerin. **(C)** MA plot for the 15,323 union pS22-Lamin A/C-binding sites, highlighting gained and lost pS22-Lamin A/C-binding sites. Sum of pS22-Lamin A/C ChIP FE scores within +/–250 bp of site center is compared. Normal, normal fibroblast GM07492. Progeria, progeria fibroblast AG11498. **(D)** pS22-Lamin A/C, c-Jun, and H3K27ac ChIP-seq profiles at the *FHL-1* locus highlighting gained pS22-Lamin A/C-binding sites. Signals are FE scores. The *FHL-1* gene is among the genes up-regulated in progeria fibroblasts, and its mutations cause cardiac and skeletal myopathies. Normal, GM07492 normal fibroblast. Progeria, AG11498 progeria fibroblast. **(E)** c-Jun ChIP-seq FE score distribution around gained pS22-Lamin A/C-binding sites and lost pS22-Lamin A/C-binding sites. **(F)** Same as **(E)** but H3K27ac ChIP-seq FE scores are shown. **(G)** Number and fraction of genes linked to gained, lost, and steady pS22-Lamin A/C-binding sites. **(H)** DisGeNet-curated disease terms over-represented among progeria-down genes linked or not linked to lost pS22-Lamin A/C-binding sites.

